# A wave of embryonic bipotent T/lymphoid tissue inducer progenitors regulates the maturation of medullary thymic epithelial cells

**DOI:** 10.1101/791103

**Authors:** Ramy Elsaid, Sylvain Meunier, Odile Burlen-Defranoux, Francisca Soares-da-Silva, Thibaut Perchet, Lorea Iturri, Laina Freyer, Paulo Vieira, Pablo Pereira, Rachel Golub, Antonio Bandeira, Elisa Gomez Perdiguero, Ana Cumano

**Author notes:** Correspondence should be addressed to A.C.

## Abstract

Multiple waves of hematopoietic progenitors with distinct lineage potentials are differentially regulated in time and space. We show that the first thymic seeding progenitors comprise a unique population of bipotent cells that generate lymphoid tissue inducer and invariant V*γ*5^+^ T cells. Both populations are of embryonic origin and induce the maturation of medullary thymic epithelial cells. Indeed, temporal depletion of the first wave of thymocytes results in a five-fold reduction of mature medullary thymic epithelial cells, after birth. We further show that these progenitors are of hematopoietic stem cell, and not, of yolk sac origin, despite the temporal overlap between the onset of lymphopoiesis and the transient expression of lymphoid transcripts in yolk sac precursors, that does not impact their strict erythro-myeloid potential. Our work highlights the relevance of the developmental timing on the emergence of different lymphoid subsets required for the establishment of a functionally diverse immune system.

## INTRODUCTION

During embryogenesis, the thymus is colonized by two waves of hematopoietic progenitors with distinct potentials, named early thymic progenitors (ETPs)^1^. The first wave of ETPs follows a unique developmental program as they share phenotype, transcriptional signature and restricted T cell differentiation potential with HSA^lo^ α4β7^-^ fetal liver (FL) common lymphoid progenitors (CLPs)^2, 3^. These CLPs are no longer found at E15.5, a stage after which the thymus is colonized by a second wave of ETPs that share phenotype, transcriptional signature and differentiation potential with lympho-myeloid primed progenitors (LMPPs)^2–5^.

Three lymphoid subsets are exclusively developed during embryonic life, they comprise T lymphocytes V*γ*5^+^ that reside in the skin, also named dendritic epidermal-like T cells (DETC), V*γ*6^+^ that reside in the lungs and in the genital urinary tract and innate lymphoid cells known as tissue inducers (LTi), a subset of group 3 innate lymphoid cells (ILC3)^1^. Both V*γ*5^+^ T cells and LTi are required for the development of medullary thymic epithelial cells (mTECs), through TNFRSF signaling, that results in the expression of the transcription factor *Aire,* a mechanism which is only conserved in prenatal stages^6–9^. *Aire* drives the expression of tissue-restricted antigens (TRA), required for negative selection of conventional T cells, and is concomitant with the expression of CD80 that is taken as a marker of mTEC maturation^8, 10^. Neonatal *Aire* expression is necessary and sufficient to induce life-long T cell tolerance^11^. LTi are also required for secondary lymphoid tissue organization^12, 13^ whereas V*γ*6 T cells contribute to remodeling of the tissues they colonize^14^.

Because the first ETPs that colonize the thymus were the only to generate V*γ*5^+^ T cells^3, 15^, it is important to determine whether they also generate other embryonic lymphoid cells and, importantly, the thymic LTi, that together with V*γ*5 T cells shape the thymic architecture^8^. Accordingly, the majority of embryonic thymic ILCs are ILC3, gradually lost after birth, while a small population of ILC2 is maintained throughout life^16^.

Several successive but overlapping waves of emerging hematopoietic progenitors, with different lineage potential, are differentially regulated in time and space^17, 18^. Hematopoietic cells first appear in the extra-embryonic yolk sac (YS) blood islands at around embryonic day 7-7.5 (E7-7.5), where primitive erythropoiesis occurs prior to the emergence of multi-lineage erythro-myeloid progenitors (EMPs)^19, 20^. The emergence of pre-HSCs starts at E9.5 in the aorta-gonad-mesonephros (AGM) and major arteries^21–24^, these cells colonize the FL around E10.5 and later (around E16) migrate to the bone marrow (BM)^17, 25^. HSC are distinguished from all other progenitors by their ability to long-term repopulate all hematopoietic lineages^17, 26^. Emerging HSC cannot however repopulate adult normal recipient animals and are designated as pre-HSC^26, 27^.

Most hematopoietic cells are constantly differentiating from HSCs although a few lineages are HSC independent and exclusively produced during embryonic development. For example, YS cells that contribute to erythropoiesis and megakaryopoiesis are also the source of tissue resident macrophages that persist throughout life^19, 28–30^. This layered organization of the hematopoietic system lead to the possibility that, similar to tissue macrophages, the first innate-like lymphocytes could also be YS derived. In line with this view, cells identified as lympho-myeloid restricted progenitors (LMPs) were observed in E9.5 YS before HSC activity is detected^31^. They were shown to express lymphoid associated genes (*Il7r*, *Rag2*, *Rag1*) and were proposed as the origin of the first wave of ETPs^32^. Independent reports have also converged to support the notion that V*γ*5^+^ and another *γ*δ T cell subset, the V*γ*6^+^ IL-17 producing cells, might originate from YS progenitors, independent of HSCs^33, 34^. In contrast to the above reports, V*γ*5^+^ and B1a B cells were shown to be preferentially derived from a particular HSC-like subset, transiently found in the FL but not in the adult BM, marked by an history of *Flk2* expression^35^

In this report, we undertook a characterization of the functional properties and fate of the progeny of the first ETPs as well as their developmental origin. Unlike the current view that during embryogenesis ILCs are derived exclusively from the common α4β7^+^ ILCs progenitors^3637^ we showed, in single cell differentiation assays, that the first wave of ETPs comprises bipotent T/LTi progenitors that contribute to the thymic LTi. Colonization of Rag*γ*c thymic lobes with ETPs from the first, but not from the second wave, reconstituted the thymic LTi compartment. Depletion of the first wave of lymphoid progenitors in the embryo resulted in lower numbers of V*γ*5^+^ DETC, LTi and severely reduced numbers of CD80^+^ mTEC, after birth. The first ETP-derived conventional T cells that lack TdT expression and exhibit a restricted T cell repertoire are rapidly overridden by subsequent waves of T cells^1, 38^. Therefore, the major role of the first wave of ETPs might be restricted to provide cells that drive thymic organogenesis and homeostasis^1, 10, 39^.

We further demonstrated, using an inducible lineage tracer model, where only YS derived progenitors are labeled that that the embryonic ETPs are not YS-derived but rather originate from intra-embryonic HSCs. YS-derived progenitors although showing a history of expression of lymphoid associated genes such as *Il7r*, *Rag2* and *Rag1*, did not differentiate into any lymphoid subset in vivo or in vitro. This transient expression, also found in other FL myeloid progenitors, is restricted to embryonic hematopoiesis and uncoupled from differentiation potential. Altogether our data highlight the impact of embryonic developmental timing on lymphocyte production, gene expression and the heterogeneity of the immune system.

## RESULTS

### E13 ETPs retain T/LTi lineage potential in vitro and generate LTi cells in a fetal thymic microenvironment

A detailed analysis of the transcriptional profiles of ETPs from the first (E13) and the second (E18) waves identified the overexpression, in the former, of ILC related transcripts from which stood out LTi-associated lineage (*Rorc*, *Cd4*, *Cxcr5*, *Il1r1* and *Ltb*) transcripts^3, 40^ (Sup Fig 1a). This comparison also showed that genes involved in *γ*δ T cells development (*Bhlhe40*, *Cited4*, *B4galnt2*, *Tgm2* and *Txk*) are differentially expressed between E13 and E18 ETPs (Sup Fig 1a). Of note, *Sell* and *Ly6a* were upregulated in E18 ETPs, in line with their resemblance to LMPPs^2, 3^.

In addition, the E14 DN3 thymocytes were devoid of TdT activity (Sup Fig 1b) giving rise to T cells with a restricted T cell repertoire raising the possibility that they do not contribute to the adaptive T cell, but rather to the innate-like T cell and ILC compartments.

We performed a two-step culture^2, 5^ (Fig 1a) that allowed detecting the potential of single ETPs (sorted as in Ramond et al^3^) to generate all major lymphoid lineages (T, B, ILC, NK) and myeloid cells (GM) (Fig 1b).

**Figure 1.**
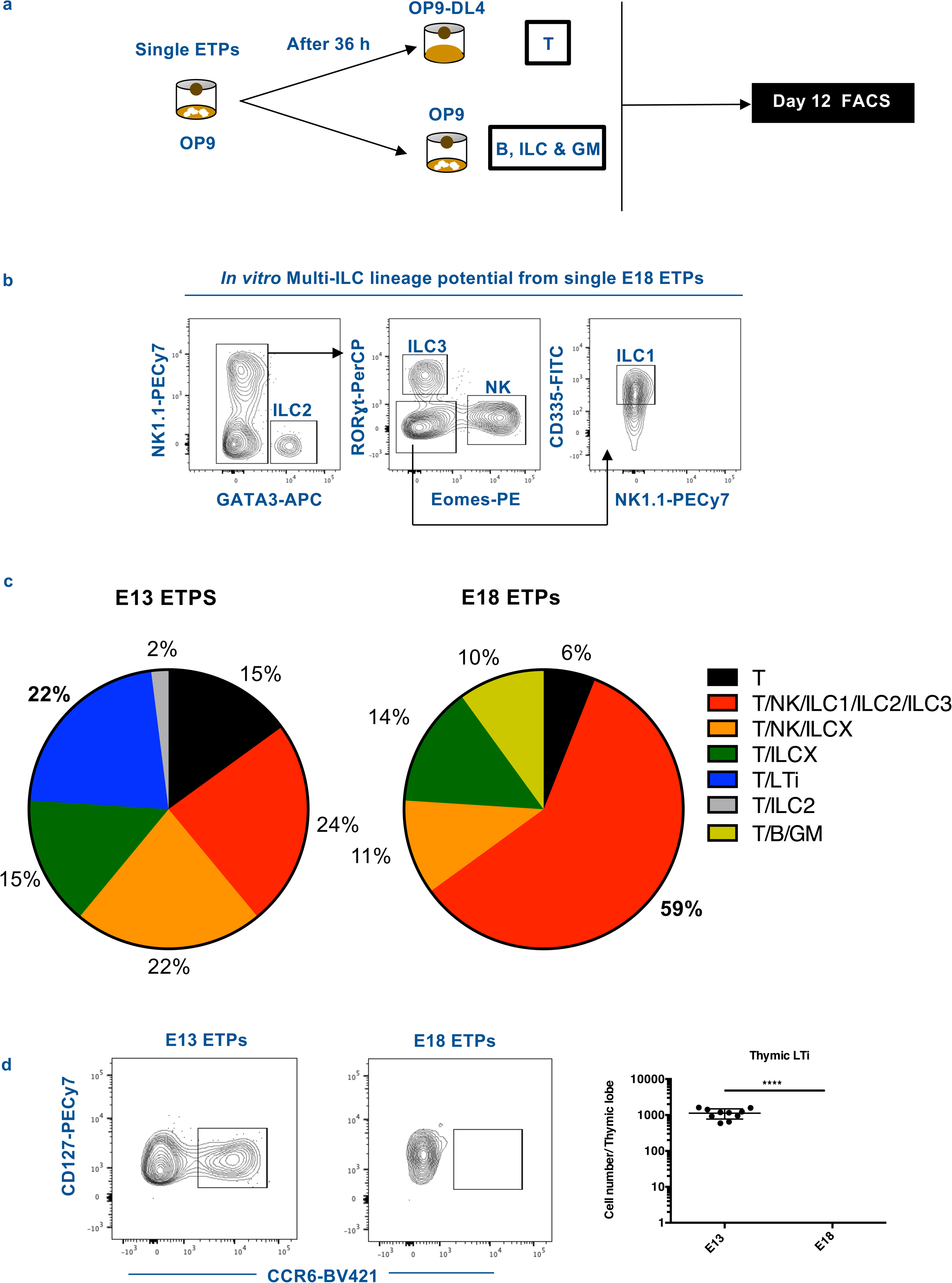
E13 ETPs retain T/LTi lineage potential in vitro and generate LTi cells in a fetal thymic microenvironment. (a) Experimental strategy for combined T, B and ILC lineage potential analysis of single ETPs. Single ETPs from either E13 or E18 were sorted onto OP9 stromal cells supplemented with IL-7, Flt3L, KitL and IL-2. After 36 hours, clones were subdivided in conditions that sustain differentiation of T cells (OP9-DL4 supplemented with IL-7, Flt3L, KitL and IL-2) or B, myeloid and ILC/LTi (OP9 supplemented with IL-7, Flt3L, KitL, IL-2 and GM-CSF). (b) Representative flow cytometry plots outlining the gating strategy used to identify ILC1, ILC2, ILC3 and NK cells of E18 ETPs single cell-derived multi-ILC lineage clone. (c) Pie chart indicating all phenotypic combinations in individual clones detected after *in vitro* differentiation of E13 (81 clones analyzed out of 240 sorted cells CE 34%) or E18 ETPs (86 clones analyzed out of 240 sorted cells CE 36%) in two independent experiments. CE, cloning efficiency. Frequency of clones generating only T cells (black); T, NK, ILC1, ILC2 and ILC3 (red) (different in E13 and E18 clones p<0.0001); T and LTi (RoRc^+^ CCR6^+^) (blue) (different in E13 and E18 clones p<0.0001); T, NK, LCX two other ILC subsets (orange); T and ILC2 (grey); T and two other ILC subsets (green); T, B and myeloid cells (olive) (different in E13 and E18 clones p=0.0036). All other combinations were not significantly different in E13 and E18 clones. (d) Flow cytometry plots of Rag2^-/-^*γ*c^-/-^ E14.5 thymic lobes reconstituted with 500 E13 ETPs or 500 E18 ETPs per lobe and gated CD3^-^ and stained for the expression of CD127 and CCR6 (left plots). Numbers of thymic LTi cells per lobe (right plot). FTOCs were analyzed at day 12 after reconstitution. **** *P* < 0.0001.

Consistent with our previous observations that ETPs before E15.5 are restricted lymphoid progenitors biased for T cell generation^3^, we found that E13 ETPs only generated T and ILC whereas E18 ETPs retained B and myeloid potential. Most ETPs gave rise to T and ILCs, although a few (6-15%) only generated T cells and none gave ILCs only. About 59% of the clones derived from E18 ETPs could generate all subsets of ILC and NK cells, whereas only 24% of E13 ETPs showed that capacity (Fig 1b, c), suggesting they are more restricted in their lineage potential.

The most striking difference between the two subsets was the high frequency of bipotent T/LTi (22%) precursors in E13 ETPs whereas no such restricted colonies were observed in the progeny of E18 ETPs (Fig 1c). Interestingly E13-derived ILC3 also expressed markers that identify LTi (CCR6^+^).

We performed fetal thymic organ culture (FTOCs), where non-irradiated Rag2^-/-^*γ*c^-/-^ thymic lobes were colonized by limited numbers of E13 or E18 ETPs, and found that only the former could generate thymic LTi (Fig 1d).

These results showed that mid gestation (E12.5-15.5) ETPs appear LTi biased and are unique in generating thymic LTi in addition to V*γ*5^+^ DETC^3^ within a thymic microenvironment.

### First wave ETPs have a LTi primed transcriptional profile

To understand the molecular basis of the LTi lineage potential differences between E13 and E18 ETPs, we performed single-cell transcriptional analysis by multiplex qRT-PCR for the expression of 41 lymphoid-associated genes. Unsupervised hierarchical clustering identified 4 distinct clusters (Fig 2a). Cluster I comprised a majority (76%) of E18 ETPs and showed no expression of ILC transcripts, indicating no ILC lineage priming consistent with their capacity to generate all ILC populations. Clusters II and IV were essentially composed of E13 ETPs and were characterized by the expression of the ILC associated transcripts *Tox*, *Tcf7* and *Gata3*. These two cluster were discriminated by the expression of the LTi cell transcripts *Cxcr6*, *Cd4*, *Rorc*, *Cxcr5*, *Il1r1* and *Ltb* in Cluster IV, consistent with a ILC3 restricted differentiation potential. Cluster II instead expressed *Zbtb16, Gzmb, Nfil3, Itga2b* and *Gzma* associated with innate-like T cells^36^. Cluster III comprised two thirds of E13 and one third of E18 ETPs and was characterized by the expression of lower levels of *Tox*, *Gata3*, *Tcf7*, *Sox4*, *Runx1* and *Runx3* indicative of common ILC priming and consistent with a broad ILC differentiation potential (Fig 2a). Taken together the results above indicated that the first wave of ETPs are primed to generate LTi and invariant T cells.

**Figure 2.**
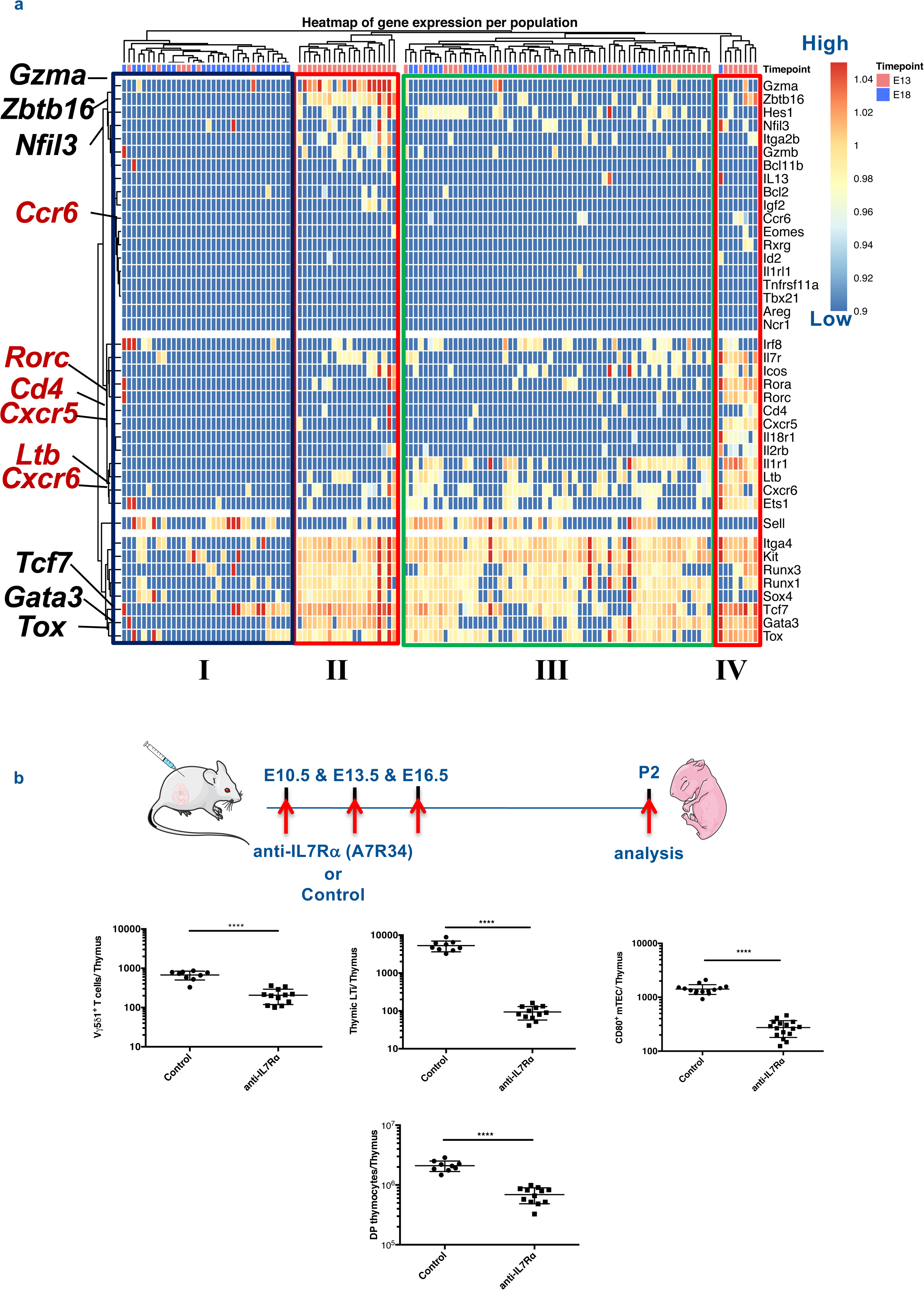
E13 ETPs have a transcriptional LTi signature and are required for the maturation (CD80^+^) of mTEC. (a) Single-cell multiplex qPCR analyzed by hierarchical clustering of single E13 and E18 ETPs (80 cells each) for the expression of ILC lineage specific transcripts (right margin). Each column represents a cell (E13 ETP in salmon, E18 ETP in blue) and each row represents one gene (of 41 genes). Highlighted are ILCs associated transcripts (black) and LTi associated transcripts (in red). Analysis was done by normalizing expression to two independent house-keeping genes (*Gapdh* and *Actinb*) in R package Phenograph as in (Perchet et al., 2018) and represented in a code color where red represents high expression and blue low expression. Data are pooled from two independent experiments. (b) Pregnant females were injected at E10.5, E13.5 and E16.5 with 150μg of IL-7Ra antibody A7R34. Thymic lobes were analyzed at P2 for the presence of V*γ*5Vδ1 *γ*δT cells, for LTi (CD127^+^ CCR6^+^ TcR^-^ CD3^-^ cells) and for CD80^+^ mTEC (EpCAM^+^ Ly51^-^ UEA.1^+^ CD45^-^). Plots show numbers of the respective populations per thymus. Data from two independent experiments. **** *P* < 0.0001.

### First wave ETPs contribute to the maturation of the thymic mTEC

LTi and invariant *γ*δ T cells have been shown to induce maturation of thymic mTEC^10^. To directly assess the role of the first wave ETP in mTEc maturation, we depleted IL-7Ra expressing FL cells, that are the precursors of the first wave ETPs with anti-IL7Rα antibody injected in pregnant females. It has been previously shown that A7R34 has a depleting effect in IL-7Rα expressing cells^41–43^ and by giving three consecutive injections (at E10.5, E13.5 and E16.5 of gestation) we expected to severely deplete the first wave of ETPs and their progeny. Thymus of treated mice analyzed at P2 had significantly lower numbers of V*γ*5^+^ T, CD4^+^CD8^+^ double positive (DP) cells and less than 10-fold LTi, compared to controls, indicating that the first wave ETPs and their progeny had been efficiently depleted (Fig 2b). Interestingly, mTEC expressing CD80 (a surrogate of *Aire* expression)^8^ were more than 5-fold reduced indicating that the first wave ETPs is required for the maturation of thymic mTEC. Taken together our results indicate that the first wave ETPs contribute to the maturation of mTEC that ensure conventional T cell negative selection.

### Neonatal thymectomy has no effect on tissue colonization by embryonic derived Vγ5**^+^** and Vγ6^+^ γδ T cells

V*γ*5^+^ and V*γ*6^+^ *γ*δ T cells in the skin and in the lymph nodes (LN), respectively, are of embryonic origin^15, 44^. To assess the dependency of these embryonic tissue resident *γ*δ T subsets on thymic output that might inform on their origin in the first or second wave of ETPs, we performed neonatal thymectomy. Newborn (up to 36 hours after birth) mice were thymectomized or sham-operated as a control (Fig 3a). At 6 weeks of age complete thymectomy was ascertained by the absence of double positive CD4/CD8 cells in the tissue around the thymus location (Fig 3b). Thymectomized mice (n=6) were severely lymphopenic as shown by the numbers of CD4 and CD8 αβ T cells in the inguinal (i) lymph node (LN) (Fig 4d) and in the spleen; of note, CD44^-^ αβ T cells in the spleen were severely reduced (Fig 3e), a sign of activation of the T cell compartment consequent to the lymphopenia and consistent with the absence of a functional thymus in these mice. The numbers of V*γ*5 ^+^ *γ*δ T cells in the skin and of iLN V*γ*4^-^ IL17 producing *γ*δ T cells (considered as V*γ*6^+^ *γ*δ T) were similar in sham and in thymectomized animals (Fig 3c, d). By contrast, iLN V*γ*4^+^ IL-17 producing *γ*δ T cells were severely reduced (Fig 3d). These experiments suggest that the two embryonic *γ*δ T cell subsets are likely derived from the first wave of ETPs. All other αβ or *γ*δ T depend on post-natal thymic output and thus are likely largely issued from the second wave of ETPs.

**Figure 3.**
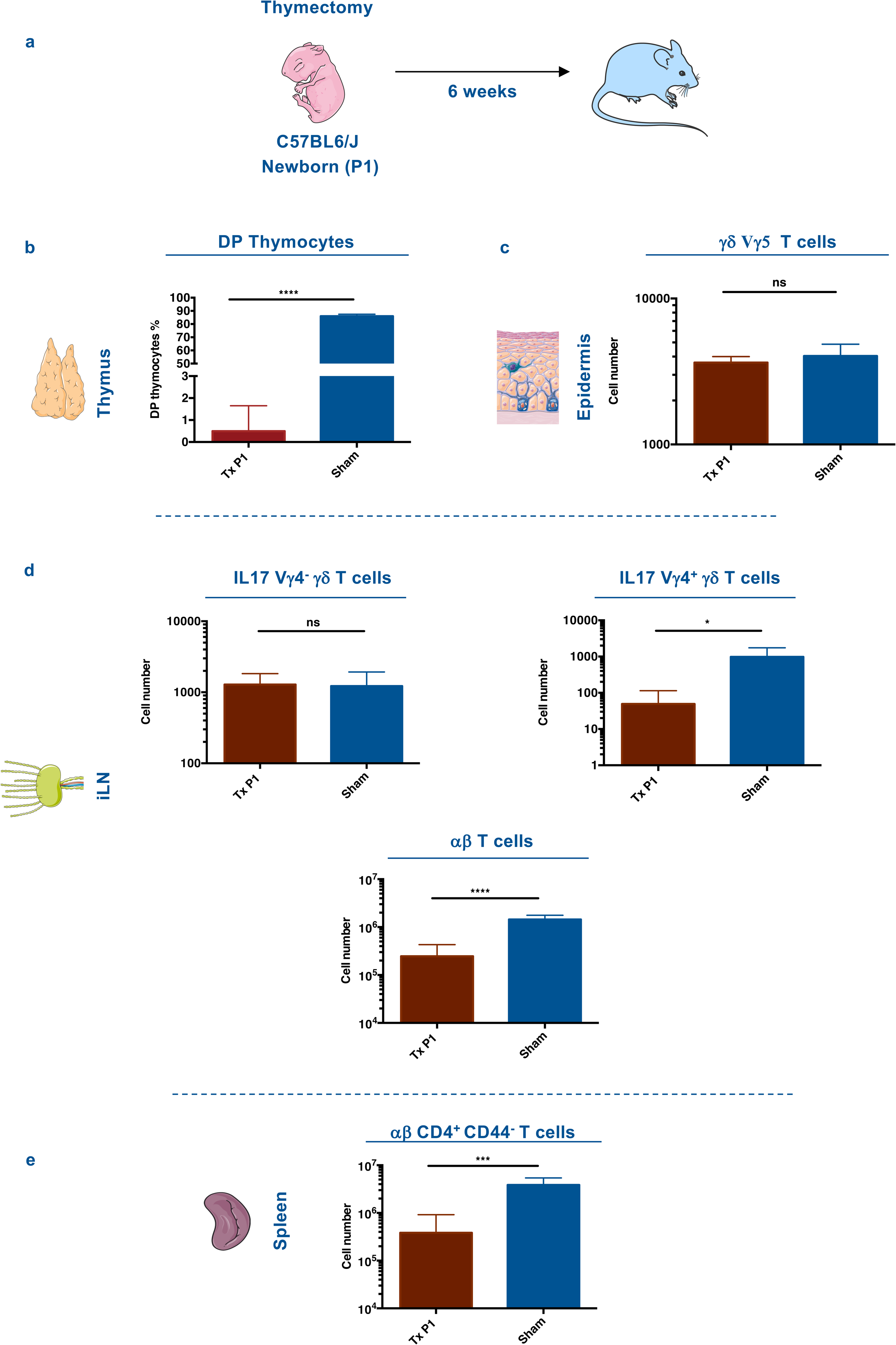
Neonatal thymectomy has no effect on embryonic derived Vγ5 and Vγ6 γδ T cells. (a) Schematic diagram of neonatal thymectomy experiment. P2 newborn mice were thymectomized and analyzed 6 weeks later. (b) Frequencies of double positive CD4/CD8 T cells found in the thymus and peri thymic tissue isolated from sham and thymectomized mice, respectively. (c) Number of V*γ*5^+^ *γ*δ T cells in the epidermis isolated from one ear of sham and thymectomized mice. (d) Number of V*γ*6 IL17 *γ*δ T cells, V*γ*4 IL17 *γ*δ T cells and αβ T cells in one inguinal lymph node, isolated from sham and thymectomized mice. (e) Number of αβ CD4^+^ CD44^-^ T cells in the spleen. \**P* < 0.05, \*\**P* < 0.01 and \*\*\**P* < 0.001 (Student’s *t*-test). Data are representative of at least 4 mice in each group from two independent experiments. Data are depicted as mean ± s.e.m.

### Embryonic ETPs have multi-ILC lineage potential *in vivo*

We assessed the ILC lineage potential of E13 and E18 ETPs *in vivo.* CD45.2^+^ E13 or E18 ETPs were intravenously transferred into 1-day old Rag2^-/-^γc^-/-^ CD45.1^+^ mice. Recipient mice were analyzed at 5 weeks post-transfer^45^ (Sup Fig 2a). While mice engrafted with E13 ETPs only generated ILC, E18 ETPs also generated B and myeloid cells, (Sup Fig 2b) consistent with the more restricted differentiation potential of E13 ETPs. Analysis of PB 3 and 6 weeks or splenic cells 5 weeks, after transplantation show a total absence of B and myeloid progeny from E13 ETPs (Sup Fig 2b, e). As expected, few T cells were generated in these chimeras because of the atrophic thymus in Rag2^-/-^γc^-/-^ mice and because TSP down-regulate CCR7 and CCR9 as they enter the thymus and consequently lose their ability to home back to the thymus^46^. Analysis of intestinal lamina propria, where all ILC lineages coexist, showed that ETPs from both waves generated NK cells (EOMES^+^), ILC1 (EOMES^-^), ILC3 (RORɣt^+^) and ILC2 (GATA3^+^). In the lung we found ILC2, in the spleen NK cells and in the liver NK cells and ILC1 (Sup Fig 2c, d). These results indicated that E13 and E18 ETPs have the capacity to generate all ILC subsets *in vivo* recapitulating the expected tissue distribution.

### Emergence of *Il7r* expressing YS-progenitors in E9.5 embryos

YS derived progenitors have been shown to express lymphoid associated genes such as *Rag*, *Flt3* and *Il7r*^31, 32^, suggesting the possibility that the first ETPs originated in an HSC independent pathway. This notion was further reinforced by a recent report suggesting a YS origin of V*γ*5 ^+^ *gd* T cells^33^.

To identify where and when the first *Il7r*-expressing cells can be detected we used a lineage tracer mouse line, *Il7rα*^Cre^Rosa26^YFP 47^. We performed time course analyses of YFP expressing cells in the YS, AGM region and FL (Fig 4a). No YFP expression was observed in E8.5 embryos, in any of the analyzed tissues in either the Kit^+^ or Kit^-^ fraction (data not shown). Consistent with previous studies, YFP^+^ progenitors were first detected in Lin^-^Kit^+^CD41^+^ E9.5 YS (Fig 4b, c) a stage at which they were undetected in the AGM or in the placenta (Fig 4d). 5-8 % of the c-kit^+^ YS cells were YFP^+^ (Fig 4b, c) expressed CD31, CD16/32 and lacked surface expression of CD115, CD127, CD135, the phenotype of YS EMPs (Lin^-^Kit^+^CD41^+^YFP^-^)(Fig 4e)^19, 30^.

**Figure 4.**
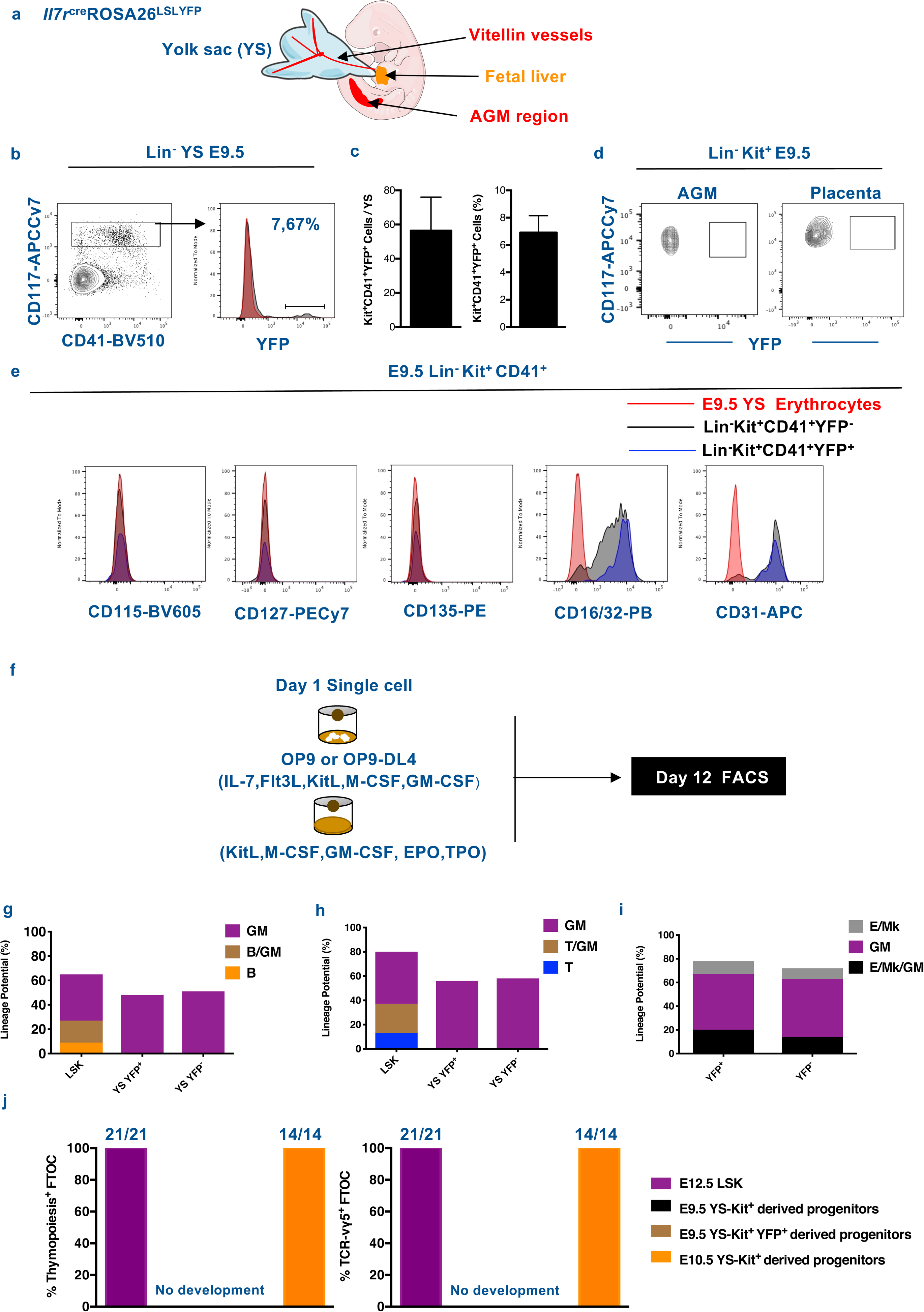
Emergence of *Il7Rα^+^* expressing YS-progenitors in E9.5 embryos. (a) Schematic representation of the anatomical sites analyzed in mouse embryos: YS, head, fetal liver and the AGM region. (b) Representative flow cytometry plot showing YFP expression on live Lin^-^ CD117^+^ CD41^+^ E9.5 YS cells from *Il7r*^cre^ROSA26^LSLYFP^. (c) Number and Percentage of YFP^+^ per YS (mean ± s.e.m). Data representative of 14 embryos from 4 independent experiments. (d) Representative flow cytometry plot showing YFP expression on live Lin^-^ CD117^+^ E9.5 placenta and AGM region cells. (e) Expression of CD115, CD127, CD135, CD16/32 and CD31 within the E9.5 YS Lin^-^ CD117^+^ CD41^+^ YFP^+^ and YFP^-^ populations. (f) Experimental strategy for single cell lineage potential assay of E9.5 YS Lin^-^ CD117^+^ CD41^+^ YFP^+^ and YFP^-^ populations. (g) Frequency of wells containing B cells or (h) T cells in cultures of single sorted E9.5 YS Lin^-^ CD117^+^ CD41^+^ YFP^+^ and YFP^-^ cells (180 cells from each population analyzed in three independent experiments). (i) Frequency of erythroid and megakaryocytic (E/Mk), myeloid (GM) or mixed E/Mk/GM colonies of single sorted E9.5 YS Lin^-^ CD117^+^ CD41^+^ YFP^+^ and YFP^-^ cells. (j) Frequency of thymic lobes irradiated and reconstituted with CD117^+^ E9.5 YS YFP^+^ or YFP^-^, or with E12.5 LSK or E10.5 YS as controls, that contained CD4/CD8 DP cells (left plot) or TCR-Vγ5^+^ T cells (right plot). FTOCs were analyzed at day 12 after reconstitution.

To investigate the potential of YS YFP^+^ cells to develop into T/GM and B/GM cells, single cells were cultured into monolayers of OP9-DL4 or OP9 stroma (Fig 4f). While YFP^+^ YS cells showed myeloid potential, they lacked detectable T or B cell potential (Fig 4g, h). Liquid-culture colony assays showed that both YS Lin^-^Kit^+^YFP^+^ and Lin^-^Kit^+^YFP^-^ cells exhibited comparable GM and MkE potential (Fig 4i). These results indicated that expression of YFP in the E9.5 YS embryos does not equate with lymphoid potential, and that both YFP^+^ and YFP^-^ E9.5 YS Kit^+^ cells showed a lineage potential similar to EMPs^19, 28, 30^.

To determine whether E9.5 YS Kit^+^ progenitors could give rise to the first differentiating *γ*δ T cells, the V*γ*5^+^ T cells, we performed FTOCs of thymic lobes colonized with either E9.5 YS Kit^+^ YFP^+^ progenitors, E10.5 YS-Kit^+^ progenitors or E12 FL LSKs, as positive controls (Fig 4j). Consistent with their inability to generate T cells in OP9-DL4 cultures, E9.5 YS-derived cells did not generate V*γ*5^+^ T cells or initiate thymopoiesis. By contrast, E10.5 YS-Kit^+^ progenitors and E12 FL LSK readily generated T cells including V*γ*5^+^ T cells. E10.5 YS-Kit^+^ progenitor derived T cells originate from AGM-derived pre-HSC, known to enter circulation between E9.5 and E11.5^48, 49^ and were used to probe the sensitivity of the T cell assay.

These experiments showed that E9.5 YS progenitors are devoid of lymphoid potential under culture conditions sufficiently sensitive to detect few circulating T cell progenitors.

### Expression of lymphoid associated transcripts in YS progenitors is transient

To further characterize YFP^+^ YS progenitors that share phenotype and differentiation potential with EMPs, we analyzed the expression of transcripts associated with multiple hematopoietic lineages in single E9.5 YS YFP^+^ and YFP^-^ Lin^-^CD117^+^CD41^+^ cells. Unsupervised hierarchical clustering did not segregate YFP^+^ and YFP^-^ cells indicating that they do not differ significantly in the expression of the analyzed transcripts (Fig 5a). Cluster I associated *Pu-1*, *Irf8* and *Csf1r* with low frequency of *Kdr*, *Tek*, *Runx1*, *Mpl* and *Foxo1* expressing cells, a pattern of expression corresponding to EMPs^50^ and did not express *IL7ra* or *Flk2* transcripts (Fig 5a). Cluster II expressed *Pu-1*, *Irf8* and *Csf1r* and high frequency of *Kdr*, *Tek*, *Runx1*, *Mpl* and *Foxo1* suggesting that they were EMP recently specified from hemogenic endothelium^50–52^. These cells exhibited *Il7ra* and *Flk2*. Clusters III and IV comprised YFP^+^ and YFP^-^ cells with high levels of *Gata1* indicating their erythroid lineage engagement. Cells in cluster IV also expressed *Klf1* indicating that they are more advanced into the erythroid lineage differentiation. As expected, no lymphoid associated transcripts were detected in cells from cluster III and IV. *Ebf1*, *Pax5* or *Rag1* were not detected in any of the analyzed YS cells. Within the 47 YFP^+^ cells analyzed only 6 co-expressed *Il7ra* transcripts reinforcing the notion that *Il7ra* expression is transient in E9.5 YS progenitors (Fig 5b).

**Figure 5.**
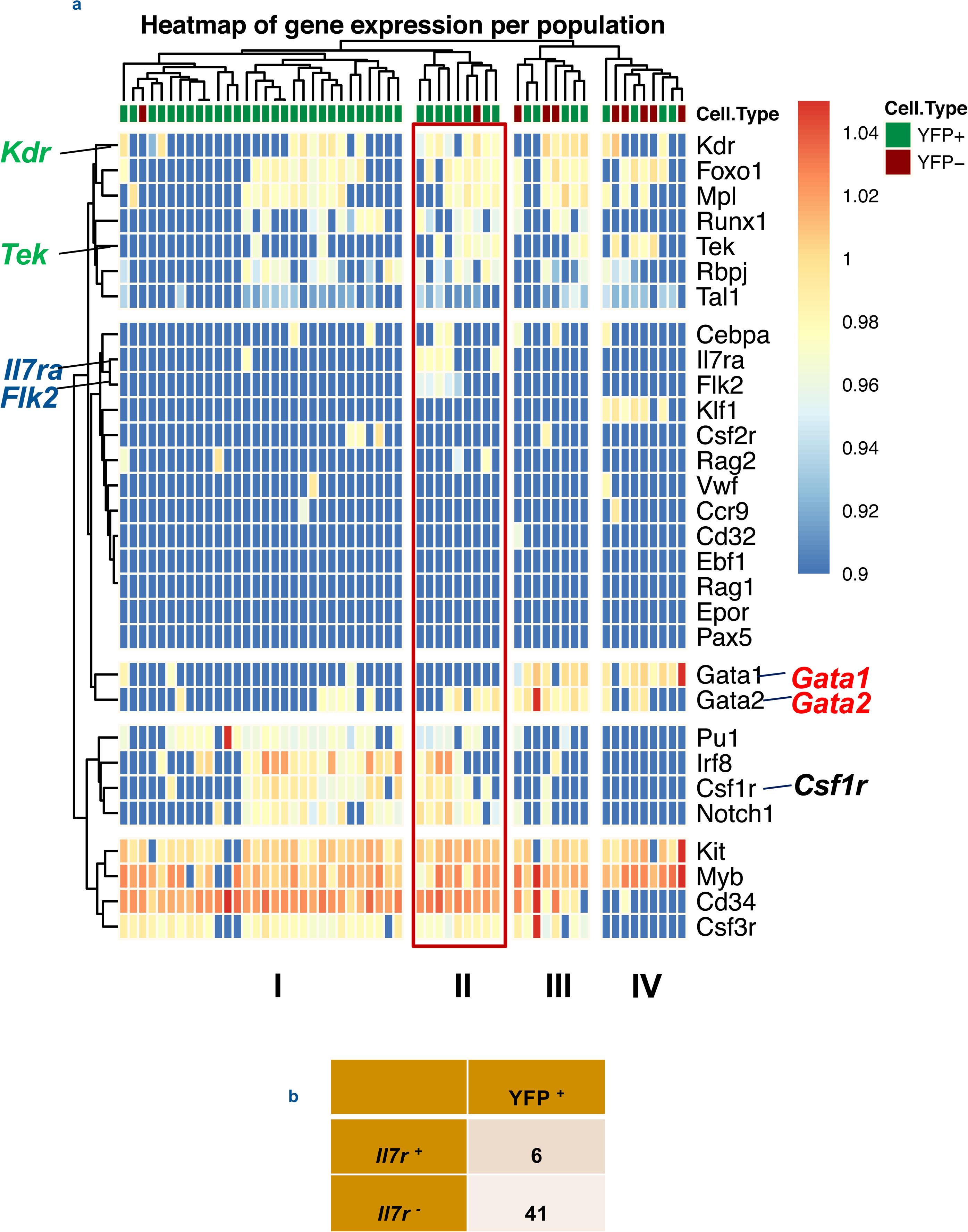
Gene expression analysis of single E9.5 YS Lin^-^ CD117^+^ CD41^+^ YFP^+^ and YFP^-^ cells. (a) Single-cell multiplex qPCR analyzed by hierarchical clustering of single YFP^+^ (47 cells) and YFP^-^ (10 cells) E9.5 Lin^-^ CD117^+^ CD41^+^ cells analyzed for the expression of lymphoid associated (*Il7r*, *Rag1*, *Rag2*, *Flt3*), myeloid associated (*Cebpa*, *Cd32*, *Csf1r*, *Csf2r*, *Csf3r*, *Irf8*, *Pu1*), B cell associated (*Pax5*, *Ebf1*), E/Mk associated (*EpoR*, *Gata1*, *Mpl*, *Tal1*, *Vwf*, *Klf1*) and hematopoiesis associated (*Gata2*, *Myb*, *Runx1*, *Kit*, *Cd34*) genes. (b) Number of *Il7r^+^* and *Il7r^-^* cells within E9.5 YS YFP expressing cells.

Interestingly, analyzed at the population level, YFP^+^ cells expressed significantly higher levels of *Rag1*, *Rag2* and *Il7r* than YFP^-^ cells albeit 10-fold lower than CLP (Sup Fig 3).

These experiments indicated that lymphoid associated gene expression, only detected in emerging YS-derived EMP, is transient and as shown previously, dissociated from differentiation potential.

### Multiple FL hematopoietic progenitors express lymphoid associated genes

The previous results dissociated the expression of YFP from lymphoid potential in *Il7ra*^Cre^Rosa26^YFP^ embryos. We further assessed whether similar expression also occurs in other embryonic hematopoietic cells. We found that unlike in adult BM where *Il7r*^Cre^ labels lymphoid progenitors^47^, YFP expression in the FL was found in multipotent and myeloid progenitors (Sup Fig 4a, b). FL and fetal blood (FB), short-term HSCs (ST-HSCs), multipotent progenitors (MPPs), granulocytes-myeloid progenitors (GMPs), common myeloid progenitors (CMPs) and megakaryocytes-erythroid progenitors (MEPs) were labeled at frequencies ranging from 10%-40% (Sup Fig 4a, b; see Sup Figure 5 for gating strategy). The frequency of YFP^+^ ST-HSC and myeloid progenitors progressively decreased after birth and was undetectable in adult mice^47^ (Sup Fig 4b). FL YFP^+^ GMPs and CMPs lacked any detectable B cell potential *in vitro*, retaining GM potential (Sup Fig 4c).

To establish whether YFP^+^ cells contribute to other blood lineages we analyzed the brain microglia, the FL Kupffer cells and the skin mast and Langerhans cells of E18 *Il7ra*^Cre^Rosa26^YFP^. We found expression of YFP in about 40% of all these populations (Sup Fig 6a, b). These observations indicated that at early stages of development *Il7ra*^Cre^ also labels cells devoid of lymphoid potential, and therefore this model does not faithfully trace the origin of embryonic lymphoid cells.

### Thymopoiesis-initiating cells develop exclusively from HSC-derived progenitors

To define the relative contribution of YS cells and HSCs to the first wave of ETPs, we used a fate mapping mouse model expressing the tamoxifen inducible Cre (*MerCreMer*) under the control of the *Csf1r* promoter^53^ (Fig 6a). Injection of hydroxy-tamoxifen (4-OH-TAM) in either *Csf1r*^MeriCreMer^*Rosa26*^YFP^ or *Rosa26*^Tomato^ (depending on the availability of fluorochrome-labelled antibodies for the analysis) induced recombination and permanently labelled *Csf1r* expressing cells and their progeny. *Csf1r* is first expressed in E8.5 YS progenitors^28^, later it is expressed in E9.5 YS *Il7ra*^Cre^Rosa26^YFP^ (Fig 5a, Sup Fig 3)^31^ and at E10.5 *Csf1r* is also expressed in pre-HSC^51^ (Sup Fig 7d). Consequently, single 4-OH-TAM injection at E8.5 or E9.5 should mark *Csf1r*^+^ myeloid cells and EMPs of YS origin that also comprise *Il7r* expressing cells. Injection 4-OH-TAM at E10.5 should mark HSC derived progenitors and *Csf1r*-expressing myeloid cells. In embryos pulsed at E8.5, YFP^+^ cells were detected in the E9.5 YS. These cells expressed *Csf1r*, *Il7r* and were devoid of lymphoid potential (Sup Fig 7a, b, c). In embryos pulsed at either E8.5 or E9.5 and analyzed at E12.5, YFP^+^ HSC, HSC-derived lymphoid progenitors and ETPs were either not detectable or represented less than 1% in FL and thymus, respectively (Fig 6b, c). In contrast, 30-50% of microglia cells were labeled, indicating the labeling efficiency of using this mouse model. Embryos pulsed at E10.5 and analyzed at E12.5 showed significant labeling in populations of FL HSC, multipotent, lymphoid and myeloid progenitors (Fig 6c). Interestingly, the frequency of Tomato^+^ cells in E12.5 ETPs (25%) and in FL LTi (22%) (See Sup Fig 7e for gating strategy) paralleled that of HSCs (31%).

**Figure 6.**
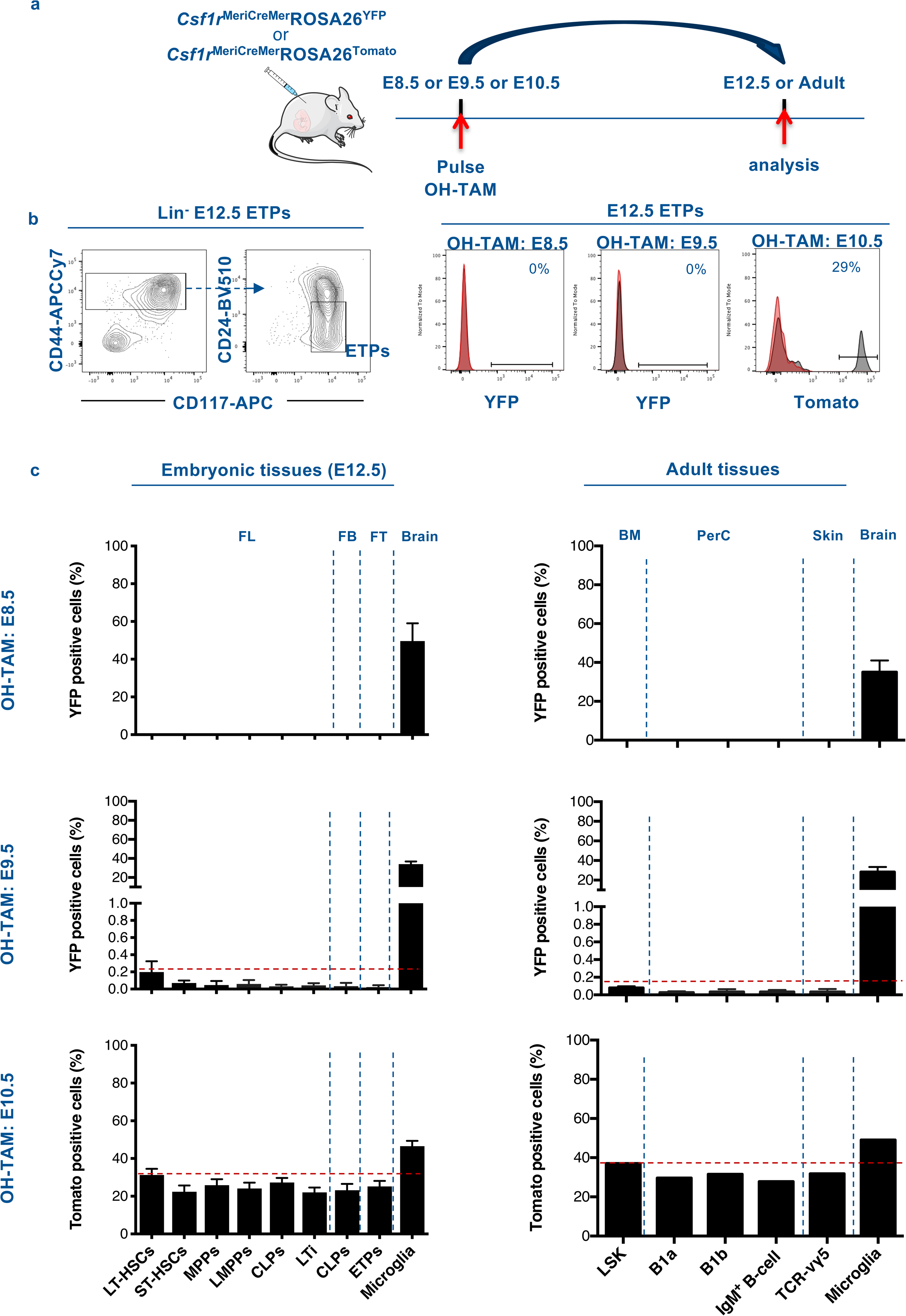
Thymopoiesis-initiating cells develop exclusively from HSC-derived progenitors. (a) Experimental design of lineage tracing analysis of *Csf1r*-expressing cells after OH tamoxifen (OH-TAM) administration at E8.5, or E9.5, or E10.5. Arrows indicate pulse and analysis time points. (b) Flow cytometry analysis of E12.5 early thymic progenitors (ETPs). E12.5 thymic lobes of embryos pulsed at indicated time points were analyzed. Left plots show the gating strategy to identify ETP. Right representative histograms showing the frequency of Tomato^+^ or YFP^+^ cells in ETP. For each experiment one representative analysis is shown. (c) Frequencies of YFP or Tomato labelled hematopoietic progenitors and Lymphoid tissue inducer (LTi) in E12.5 fetal liver, FB and thymi (left plots) and of LSK, B1 B cells, B cells and Vγ5^+^ T cells in adult tissues (right plots) of animals pulsed at E8.5, E9.5 or E10.5. Microglia served as controls for labelling efficiency. All data are pooled from minimally two independent experiments, for embryos analyzed at E12.5 (pulsed at E8.5, n = 8; E9.5, n = 10; E10.5, n = 10) and for adult tissues animals were analyzed between 10-12 weeks of age (pulsed at E8.5, n = 6; E9.5, n = 6; E10.5, n = 2). Data are depicted as mean ± s.e.m except for adult animals pulsed at E10.5 (mean). Abbreviations: ETPs, early thymic progenitors; PerC, peritoneal cavity; BM, bone marrow; FB, fetal blood; FL, fetal liver; FT, fetal thymus.

These fate-mapping experiments directly demonstrated that under physiological conditions no detectable ETPs and lymphoid progenitor originate from YS cells, rather their origin is coincident in time with that of pre-HSC.

Innate-like V*γ*5^+^ T cells or B1-a cells that differentiate during a limited time window of embryonic development have been proposed to derive from HSC-independent precursors. We analyzed V*γ*5^+^ T cells and B-1a in the skin and peritoneal cavity respectively of adult *Csf1r*^MeriCreMer^ *Rosa26*^YFP^ or *Csf1r*^MeriCreMer^ *Rosa26*^Tomato^ mice pulsed at either E8.5, E9.5 or E10.5 (See Sup Fig 7f for gating strategy).

Adult mice pulsed at E8.5 showed no YFP labelled lymphocytes and those pulsed at E9.5 showed less than 1% of V*γ*5^+^ T or B-1a labelled cells. In both cases, 30-40% of microglia cells were YFP labelled (Fig 6c). Embryos pulsed at E10.5 showed a frequency of V*γ*5^+^ T or B-1a labelled cells similar to that found in HSCs (Fig 6c). These results are consistent with a strict HSC origin of all lymphoid cells including the first wave of ETPs and innate-like lymphocytes of embryonic origin.

## DISCUSSION

Here we show that the first ETPs have a unique capacity to generate LTi that together with innate like invariant *γ*δ T cells induce the maturation of medullary thymic epithelial cells^8, 9^. A proper compartmentalization of the thymic architecture into a cortex and a medulla depends on the interaction between the first developing V*γ*5^+^ T cells, CD3^-^ LTi and the thymic epithelial cells, which is essential to generate T cell tolerance^8–10^. We have previously shown that the first ETPs derive from a unique and transient set of hematopoietic progenitors, T-biased in their differentiation potential that colonize the thymus between E12.5 and E15.5 ^3^. These precursors when compared to ETPs at later stages were the only to generate V*γ*5^+^ *γ*δ T cells^3^. We show here by single cell lineage potential assays and by colonization of a fetal thymus microenvironment, that these first ETPs comprise a population of bipotent cells that reconstitute the invariant T and LTi lineages. This observation was further supported by a transcriptional analysis showing a cluster of E13.5 ETPs that, although not committed, are primed towards the LTi lineage with high expression of *Rorc*, *Cd4*, *Cxcr5*, *Il1r1* and *Ltb*^40^. Together with the absence of detectable ILC restricted progenitors within E13.5 thymocytes and our results showing that subsequent ETPs found at later stages do not generate thymic LTi these data indicate that the first wave ETPs are unique in their functional properties. The first ETPs were also devoid of TdT activity giving rise to T cells without N sequences and thus with a restricted T cell repertoire. Low TdT expression together with the poor proliferative capacity and the observation that after P4 virtually all αβ expressing conventional T cells have N sequences in their V(D)J junctions^38^, suggested that the contribution of the first ETPs to the conventional T cell compartment is limited^1^. A recent report analyzing human embryonic thymopoiesis suggest that similar sequence of events might be shared between mouse and human^54^.

All waves of ETPs were capable of restoring the tissue resident ILC compartments of newborn Rag2*γ*c^-/-^ mice. This finding suggests that ETPs in circulation can home to the peripheral tissues and differentiate *in situ* into all ILC populations.

Our data show that the first ETPs, isolated using generally accepted surface markers^3, 55^ and not based on reporter mouse models, are T/ILC restricted with poor B and myeloid potential arguing against the notion that the embryonic thymus is initially seeded by lympho-myeloid restricted progenitors. Using *Rag1*-GFP mice, it was argued that E11.5 ETPs retained myeloid potential and resembled PIRA/B^+^ LMPPs in phenotype, lineage potential and molecular signature^32^. However, it is not clear whether these cells were ETPs because, at this stage, the epithelium that forms the thymus anlage is not folded resulting in a difficult distinction from the surrounding pharyngeal pouches. Moreover, the analysis based on the unique expression of GFP in *Rag1*-GFP embryos could include a heterogeneous population of circulating hematopoietic progenitors, thus compromising the conclusions from this study. Different strategies to isolate ETPs might explain the discrepancies in lineage potential and molecular signature between the two studies.

As the origin of progenitors during development may be linked to the generation of distinct populations within the immune system, we revisited the controversial origin of the first wave of ETPs. By analogy with the documented YS origin of tissue resident macrophages ^18, 56^, it was proposed that also early developing innate-like T cells and therefore the first ETPs were HSC independent ^31–34, 57–59^.

The evidence that YS is a potential source of early lymphocytes is based on adoptive transfer experiments, on mice that lack blood circulation or by tracing YS cells expressing lymphoid-associated genes^31, 59^.

Our analysis of YFP expressing cells under the control of the *Il7r* regulatory sequences throughout embryonic life indicated that, unlike adult hematopoietic progenitors, multipotent and myeloid restricted embryonic progenitors express lymphoid associated genes, albeit in a transient manner, without any consequent restriction in their differentiation potential. Consistent with our analysis, a recent report using the same mouse model showed not only lymphoid cells but also FL hematopoietic progenitors and tissue resident macrophages were significantly labeled^60–62^. These observations established that in hematopoiesis, gene expression does not necessarily equate with lineage potential. Collectively, those results indicated that further analysis is required before extrapolating findings obtained by the analysis of adult hematopoiesis to embryonic development.

The complexity of hematopoietic cell generation that occurs in multiple sites in overlapping time windows hampers the precise assignment of hematopoietic progenitor origin. To unambiguously assign the contribution of each anatomical site to blood cell formation, faithful lineage tracing models that trace the progeny of each source in a non-overlapping spatio-temporal manner have to be used.

The analysis of inducible lineage tracing of *Cdh5* and *Runx1* expressing cells^33^ suggested that V*γ*5^+^ T cells were generated from early emerging hematopoietic cells, precisely from YS-derived hematopoietic progenitors.

*Cdh5*, *Runx1* and *Tie2* are expressed in endothelial cells of both the YS, that produces EMPs and the AGM that is considered the source of HSC. They undergo similar processes of endothelial to hematopoietic transition (EHT)^21, 22, 24, 52^ for the generation of hematopoietic cells. Therefore, endothelial cells or their immediate hematopoietic progeny will always be labeled in both generation sites with different frequencies at different induction time points ^29, 33, 56, 63, 64^. Thus, these models do not allow a clear discrimination between progenitors originated in the YS or in the AGM. Moreover, the lack of direct comparison between the labeling efficiency in HSC and in V*γ*5^+^ T cells at different developmental stages compromise the conclusions obtained from these mouse models^33^.

*Csf1r*^MeriCreMer^ lineage tracing allowed a temporal analysis and unambiguously traced YS derived progenitors (including lymphoid-associated gene expressing progenitors) in females induced at E8.5 and E9.5, while it traced both HSC and YS-derived *Csf1r*^+^ myeloid cells, when pulsed at E10.5.

Microglia but virtually no HSC, V*γ*5^+^ *γ*δ T cells or B1a B cells were labeled by a single pulse at E8.5, also previously reported^33, 65^. In contrast, induction at E10.5 labelled HSC, V*γ*5+ T cells and B1a B cells with similar efficiencies, indicating a lineage relationship. Moreover, when pulsed at E9.5 a low (less than 1%) frequency of labelled HSC was accompanied by equally low frequency of labelled innate-like lymphocytes. These fate mapping experiments are the first *in vivo* evidence that directly demonstrate that ETPs, innate-like lymphocytes and lymphoid progenitors originate in the embryo from *Csf1r*^+^ HSCs-derived progeny, show no contribution of HSC-independent progenitors to lymphopoiesis and settle the controversy over the lymphoid ontogeny during embryogenesis. Together these data indicate that lymphopoiesis is a hallmark of HSC and HSC-derived progenitors.

The existence of restricted lymphoid progenitors in the FL between E11.5 to E15.5 suggests that the generation of embryonic-derived tissue resident lymphoid cells follows a unique developmental program^66^. These cells are produced prior to the establishment of a fully competent adaptive immunity and persist throughout life, that possibly contribute to the integrity of body barriers and act as a first line of defense against pathogens. The ability of adult ETPs to generate ILCs in vitro has been previously documented ^36, 67, 68^, however the capacity of E13 ETPs to generate thymic LTi within a thymic microenvironment is the first evidence that these cells can originate also from hematopoietic progenitors other than the classical ILCP, and in organs other than the FL where hematopoiesis occurs^17^ (Fig 7).

**Figure 7.**
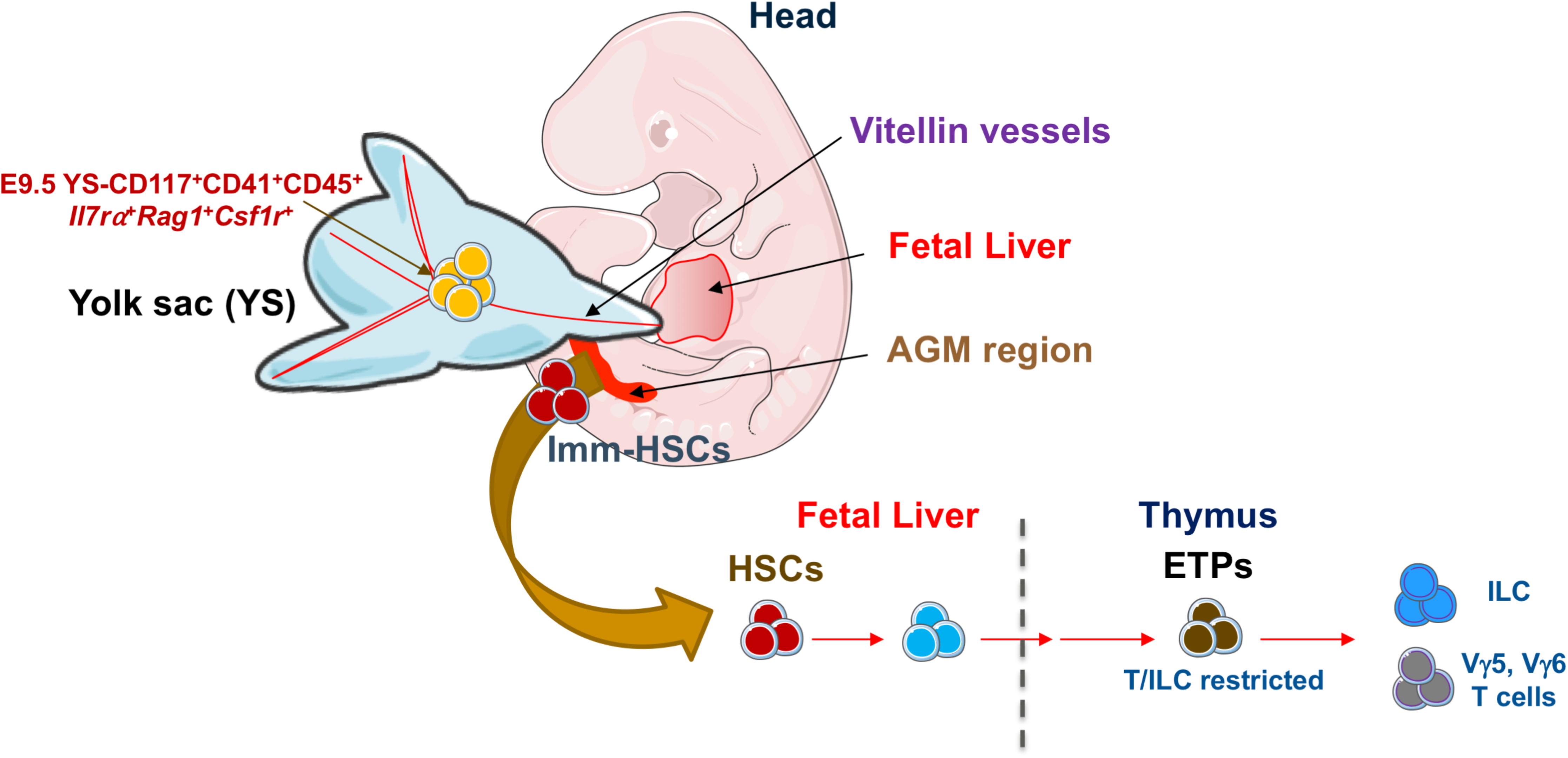
Model depicting the origin and the lineage potential of thymopoiesis-initiating progenitors. Abbreviations: ETPs, early thymic progenitors.

This highly orchestrated sequence of events highlights the requirement of a layered immune cell production whereby the first wave of ETPs is essential for thymic organogenesis and T cell tolerance.

## METHODS

### Animals

Mice were bred in dedicated facilities of the Institut Pasteur under specific pathogen-free conditions. Mattings were done overnight; on the following day, mice showing a vaginal plug were isolated and were considered to be embryonic day 0 (E0). *Il7r*^Cre 69^, Rag2^-/-^γc^-/-^ CD45.1 ^70^, *ROSA26^Tomato^* ^71^ and *ROSA26^YFP^* mice were on C57BL/6 background, *Csf1r*^MeriCreMer 53^ mice were on FVB background. All animal manipulations were performed according to the ethic charter approved by French Agriculture ministry and to the European Parliament Directive 2010/63/EU.

### Fate-mapping of *Il7r^+^* hematopoietic progenitors

For fate-mapping analysis of *Il7r+* progenitors*, Il7r*^Cre^ females were crossed to homozygous *ROSA26^YFP^*. Indicated embryonic tissues and adult tissues were analyzed by flow cytometry.

In utero pulse labelling of *Csf1r+* Progenitors. For fate-mapping analysis of *Csf1r+* progenitors, we crossed *Csf1r*^MeriCreMer^ with either *ROSA26^YFP^* or *ROSA26^Tomato^* reporter mice. For induction of reporter recombination in the offspring, a single dose of 75 µg per g (body weight) of 4-Hydroxytamoxifen (Sigma) (4OHT) supplemented with 37.5 µg per g (body weight) progesterone (Sigma) (to counteract the mixed estrogen agonist effects of 4OHT, which can result in fetal abortions) was injected either at E8.5 or E9.5 ^28^. To induce recombination in the embryos at E10.5, females received a single dose of 1.2 mg 4OHT and 0.6 mg progesterone.

In vivo anti-IL7Rα (A7R34) antibody injections. IL-7Rα blockade was performed by three successive injections of 150 ug MAb anti-IL7Rα antibody (A7R34)^72^ (a kind gift from Dr. Shin Nishikawa) i.v. in pregnant females at E10.5, E13.5 and E16.5 of gestation.

Cell suspension. Cell suspensions from embryonic, newborn and adult tissues were obtained through mechanical dissociation and/or enzymatic digestion. In brief, all cells were suspended in Hanks’ balanced-salt solution (HBSS) supplemented with 1% FCS (GIBCO). Embryonic tissues were micro dissected under a binocular magnifying lens. Organs were rinsed with HBSS plus 1% FCS to remove blood cells contamination. Then, thymus and fetal liver were passed through a 26-gauge needle of a 1-ml syringe and were filtered. Blood from embryos was obtained for 20 to 30 min HBSS supplemented with 1% FCS without calcium and magnesium. Blood cells were resuspended in a 70% (vol/vol), topped by a 40% (vol/vol), solutions of Percoll. After 20 min of centrifugation at 3000 r.p.m, cells were collected at the 40%-70% interface. For embryonic skin samples, harvested back skin was minced with scissors in RPMI medium containing 0.1 mg/ml DNase I (Roche) and 1mg/ml Collagenase I (GIBCO). Digestions were performed for 30-45 min at 37°C under continuous agitation. For embryonic head/brain, tissues were cut into small pieces, incubated in RPMI containing 10% FCS and 0.2 mg/ml Collagenase IV (GIBCO) for 30 min and then passed through a 19G needle to obtain a homogenous cell suspension. Adult thymi and spleen were dissected, and single cell suspension were obtained through a nylon mesh. Bone marrow were collected by flushing bones with HBSS. For adult skin, ears were harvested from 8-10 week-old mice, the dorsal and ventral sheets were separated using forceps and the epidermis was removed by incubation for 45 min at 37°C in 2.4 mg/ml of Dispase II (Invitrogen) and 3% FCS. Then, the epidermis was further digested for 30 min in PBS 0.1 mg/ml DNase I (Roche) and 1mg/ml Collagenase I (GIBCO). Adult brain was processed similar to embryonic brain except for an incubation period of 1 hour and cell suspension was resuspended in a 70% (vol/vol) solution of Percoll topped by a 40% (vol/vol) solution of Percoll. After 20 min of centrifugation at 3000 r.p.m, cells were collected at the 40%-70% interface.

For reconstituted Rag2*γ*c^-/-^ tissue preparations, animals were exsanguinated, and the lung was minced and incubated for 30 min at 37°C with agitation in RPMI containing 2% FCS, then incubated for 1 hour with RPMI medium containing 0.2 mg/ml Collagenase IV (GIBCO) and 0.1 mg/ml DNase I (Roche). Cells were resuspended in a 70% (vol/vol) solution of Percoll topped by a 40% (vol/vol) solution of Percoll. After 20 min of centrifugation at 3000 r.p.m, cells were collected at the 40%-70% interface. Small intestines were minced into small pieces and digested in RPMI medium containing 0.1 mg/ml DNase I (Roche) and 100 mg/ml Collagenase VIII (Sigma). Cell suspension was resuspended in a 70% (vol/vol) solution of Percoll topped by a 40% (vol/vol) solution of Percoll. After 20 min of centrifugation at 3000 r.p.m, cells were collected at the 40%-70% interface. Livers were teased and cells were resuspended in RPMI with 2% FCS, centrifuged for 7 min, then further purified after 20 min of centrifugation at 3000 in 44% Percoll. Peritoneal washouts of adult mice were performed by injecting IP PBS with 2% FCS and collecting the resulting peritoneal fluid. For neonatal thymic epithelial cells, single cell thymic suspension were enzymatically digested with Liberase TH (0.05 mg/ml; Roche) for 40 min shaking at 37 °C.

Flow cytometry and cell sorting. All digested samples were filtered, and single cell suspensions were blocked with anti-CD16/32 (BD Biosciences) and stained with antibodies listed in supplementary table 1 and fixable viability dye (Invitrogen). For transcription factor staining, cells were stained for surface markers for 30 min at 4°C then fixed according to manufacturer’s instructions (eBiosciences) and stained with fluorescent antibodies to intracellular proteins. Stained cells were analyzed on a LSR Fortessa flow cytometry or sorted using a FACSAria III (BD Biosciences). Data were analyzed with FlowJo software (Treestar).

### Culture

All experiments were done in 96-well plates at 37°C and 5% CO2 and in complete OptiMEM medium supplemented with 10% FCS, penicillin (50 units/ml), streptomycin (50 µg/ml) and β-mercaptoethanol (50 µM) (GIBCO).

**a) Frequency assay.** Single cells were sorted into 96-well plates containing a monolayer of 1000 OP9 (for B cell-ILC and myeloid potential) or OP9-DL4 (for T cell potential) stromal cells in complete medium supplemented with saturating amounts of IL-7, Flt3L, IL-2 and KitL for B, T and ILC differentiation and M-CSF, GM-CSF and KitL for myeloid differentiation. For erythroid, megakaryocytic and myeloid potential, cells were directly sorted into wells of 96-well plate and were supplemented with KitL, erythropoietin for erythrocytes differentiation, thrombopoietin for megakaryocytes differentiation, M-CSF and GM-CSF for myeloid differentiation. Cultures were supplemented with fresh cytokines every 7 d. After 12 days of culture, wells showing colonies were stained for CD19 (B cells), NK1.1 (NK cells), CD4 (T cells), CD8 (T cells), CD3 (T cells), CD11b (Myeloid cells), Gr-1(Myeloid), CD41 (megakaryocytes) and Ter119 (erythrocytes) or analyze ILC-associated transcription factors after cell fixation, GATA-3 (ILC2), RORgt (ILC3) and Eomes^-^(ILC1). Frequency scores were assigned based on the frequency of positive wells relative to the total number of plated cells.
**b) Single cell T cell-ILC-B cell-myeloid-potential assay.** Single cells were directly sorted onto a monolayer of OP9 stromal cells in Terasaki plates in medium supplemented with IL-7, IL-2, KitL and Flt3L. After 36h, growing cells were split into OP9 or OP9-DL4 cultures under the same conditions mentioned above, and clones were analyzed after 12 days. Chi-squared test was used to determine the statistically significant differences between clones obtained from E13 and E18 ETPs

### Reconstituted FTOC

Irradiated thymic lobes (30Gy) from CD45.1+ mouse embryos at E14 or E15 were colonized for 48h, on a hanging drop, in Terasaki plates by 200 YS-derived progenitors from CD45.1+ embryos at E9.5, or E10.5 or 200 LSKs from CD45.1+ embryos at E12.5, in culture medium. After colonization, thymic lobes were cultured for 12d on a filter (Millipore) floating on 3ml of complete medium in a 37°C incubator with 5% CO_2_. Then, thymi were dissociated and analyzed by flow cytometry. Non-irradiated Rag2^-/-^γc^-/-^ CD45.1 thymic lobes were colonized 500 E13 or E18 ETPs and analyzed after 12d by flow cytometry for thymic LTi.

### *In vivo* adoptive transfer

800-1200 fetal E13 HSCs, E13 and E18 ETPs from CD45.2+ embryos were sorted using FACSAria III, then injected intravenously into irradiated (1.5G) 1 day old Rag2^-/-^; γc^-/-^ CD45.1^+^ mice. Recipient mice were analyzed 5 weeks post-transfer.

### Neonatal Thymectomy

Thymectomy was performed on anesthetized mice within 36 hours of birth. The sternum was split and the bilobed thymus removed by mechanical suction and the skin sutured with surgical silk thread 8/0. After 6 weeks, the skin, the thymus, the inguinal lymph nodes and the spleen were analyzed. Controls were animals of the same age were sham operated without thymectomy.

### Quantitative RT-PCR

mRNA of sorted cells was extracted using the RNeasy Micro Kit (Qiagen), and cDNA was obtained using the PrimeScript RT Kit (Takara). qPCR was performed using TaqMan primers and TaqMan Universal Master Mix (Applied Biosystems) for the following genes: *Il7r*, *Rag1*, *Rag2*, *Csf1r*, *Csf2r*, *Csf3r* and *Myb*. Gene expression was normalized to that of *Hprt* and relative expression was calculated using the 2^-ΔCt^ method.

### Microarray

Microarray data in the GEO database with the accession code GSE50910 published by Ramond et al ^3^.

### Multiplex single-cell qPCR

Single cells were sorted in 96-well PCR plates in 9 µl of RT/pre-amp mix from the CellsDirect One-Step qRT-PCR Kit (Life Technologies) and were kept at −80°C at least overnight. For each subset analyzed, a control-well with 20 cells were sorted. Pre-amplified cDNA (20 cycles) was obtained according to manufacturer’s note and was diluted 1:5 in TE buffer, pH 8 (Ambion) for qPCR. Multiplex qPCR was performed using the microfluidics Biomark system on Biomark HD for 40 cycles (Fluidigm). TaqMan probes that were used for RT/pre-amp and qPCR are listed in supplementary table 3. Only single cells for which the three housekeeping genes could be detected before 20 cycles were included in the analysis. Analysis was done in R package Phenograph as in ^73^

### Statistical analysis

Statistical data show either mean or mean ± s.e.m., and the two tailed unpaired Student’s *t*-test was used. Statistic tests were performed using Prism software.

## ACKNOWLEDGMENTS

We thank S. Novault, S. Megharba and S. Schmutz from the flow cytometry core facility of the Institut Pasteur for technical support; the staff of the animal facility of the Institut Pasteur for mouse care. We thank H.-R. Rodewald for providing *Il7ra*^Cre^Rosa26^YFP^ mice and S.E.W. Jacobsen for fruitful discussions. This work was financed by the Institut Pasteur, INSERM, Pasteur-Weizmann Foundation and ANR (grant Twothyme) through grants to A.C.; by REVIVE (Investissement d’Avenir; ANR-10-LABX-73) to A.C. and R.E.; by recurrent funding from the Institut Pasteur, the CNRS, Revive (Investissement d’Avenir; ANR-10-LABX-73) and by an ERC investigator award (2016-StG-715320) from the European Research Council to E.G.P.

## AUTHOR CONTRIBUTIONS

R.E. designed and performed most experiments, analyzed data and wrote the manuscript; S.M. analyzed chimeric mice and contributed to discussions. F.S.S. and T.P. performed Biomark analysis. O.B.-D performed neonatal thymectomy. E.G.P., L.F. and L.I. performed fate-mapping experiments. A.B. performed adoptive transfer in neonatal mice. R.G., P.V., A.B. and P.P. contributed to the discussions. A.C. directed the research, designed experiments, analyzed data and wrote the manuscript. All authors contributed to the manuscript.

---

**Figure S1.**
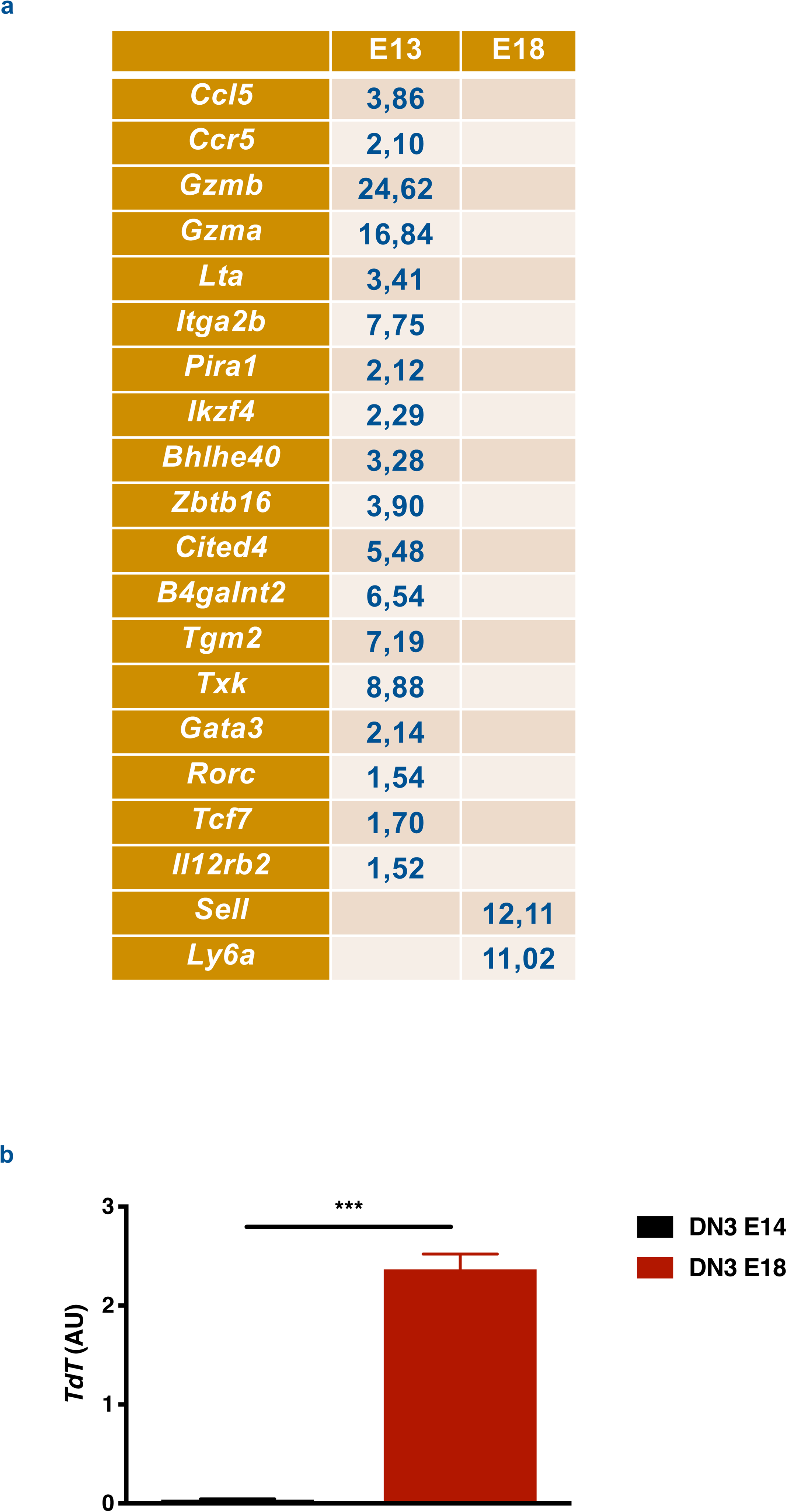
Selected genes (from the database of the Immunological Genome Project) with differential expression in TSPs at E13 versus E18. (a) Genes with differential expression in TSPs at E13 versus E18 (fold change). Transcripts analyzed are involved in either T cell development or innate lymphoid effectors development including NKT cells, ILC and γδ T cells. (b) qRT-PCR analysis of TdT expression in DN3 thymocytes at E14 versus E18.

**Figure S2.**
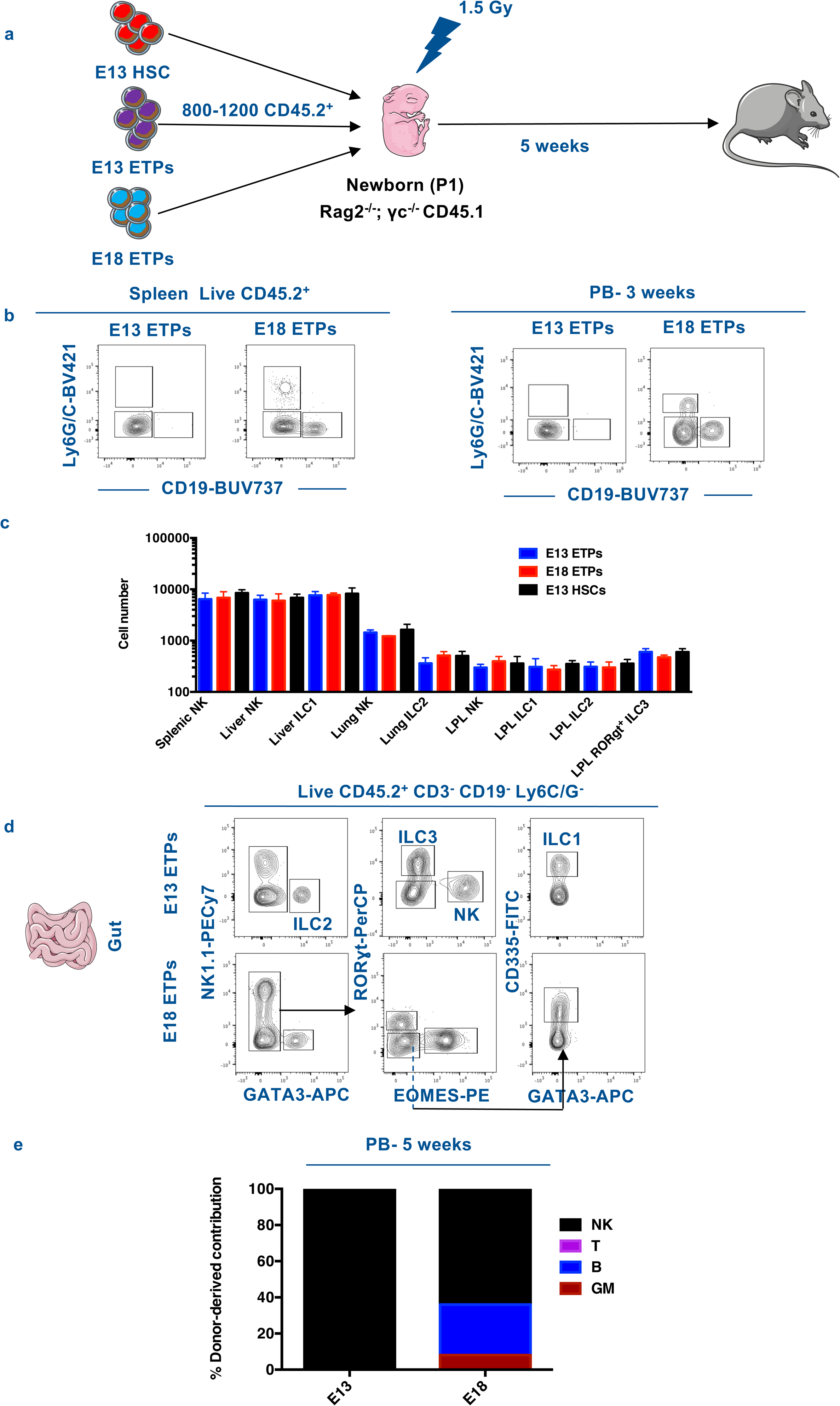
Embryonic ETPs give rise to all ILC subsets *in vivo*. (a) Schematic diagram of *in vivo* transfer experiment. P1 Rag2^-/-^γc^-/-^ mice were sub-lethally irradiated, injected with sorted E13 or E18 ETPs and analyzed 5 weeks later. (b) Flow cytometry analysis of live CD45.2^+^ splenic cells after 5 weeks and Peripheral blood (PB) after 3 weeks, for lymphoid and myeloid markers. (c) Numbers of the different ILC subsets in tissues (gut, liver, spleen and lung) of reconstituted Rag2^-/-^γc^-/-^ mice. LPL; Lamina Propria Lymphocytes. (d) Representative flow cytometry plots showing ILC analysis in the gut within viable CD45.2^+^ CD19^-^ Ly6G/C^-^ CD3^-^ cells, using the indicated surface markers and intracellular transcription factors. Data from at least 5 mice in each group from two independent experiments. (e) Frequency of donor-derived T, B, NK and myeloid lineages in the PB 5 weeks after transplantation in mice recipients of either E13 or E18 ETPs.

**Figure S3.**
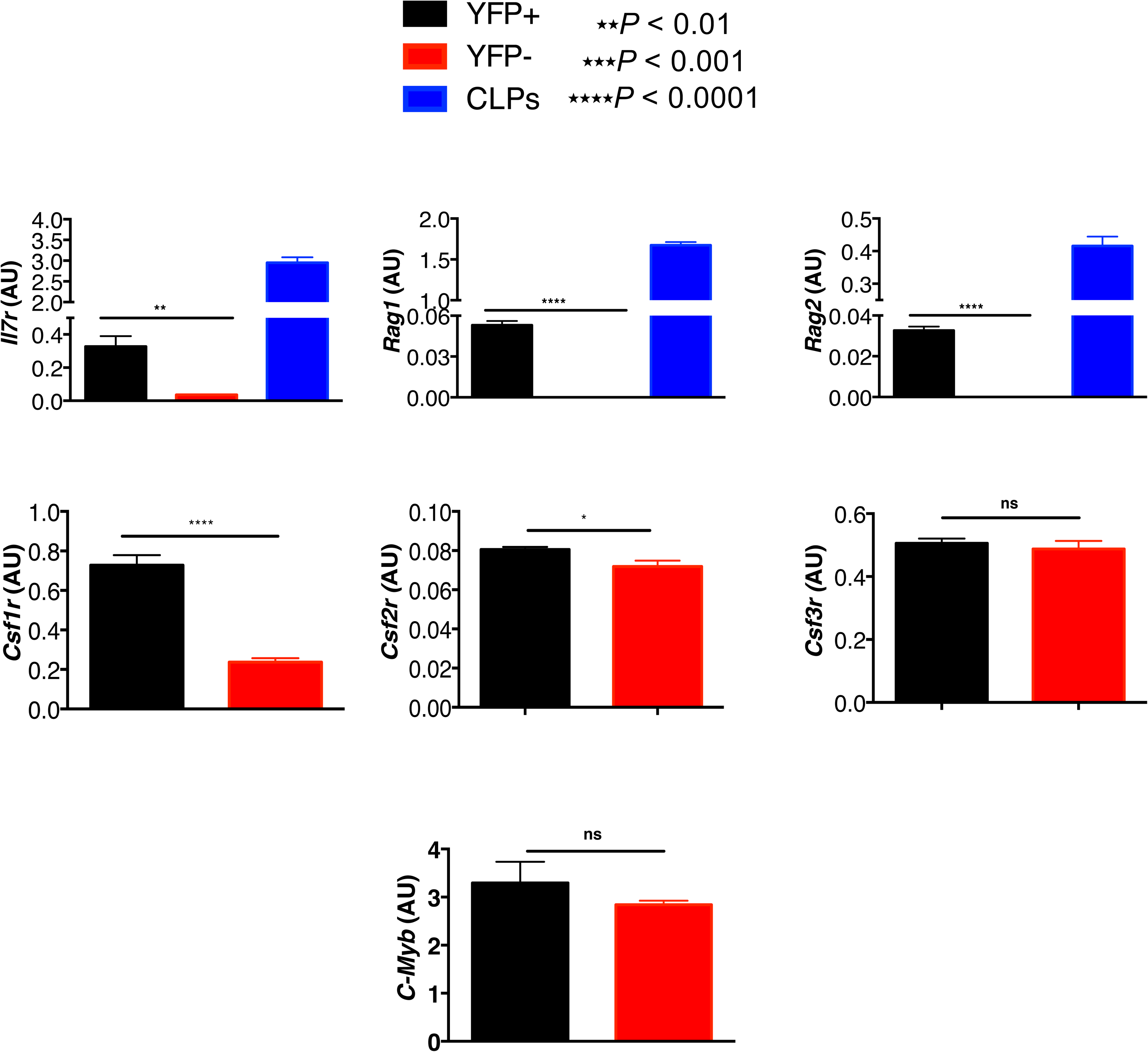
qRT-PCR analysis of lymphoid, myeloid in E9.5 YS Lin^-^ CD117^+^ CD41^+^ YFP^+^ and YFP-populations, Related to Figure 5. YFP^+^ and YFP^-^ E9.5 YS Lin^-^ CD117^+^ CD41^+^ cells were sorted and subjected to RT-PCR analysis (100 cells per reaction). Histograms show the expression of designated transcripts normalized for the expression of HPRT. CLP were taken as positive control for the expression of *il7r*, *Rag1* and *Rag2*.

**Figure S4.**
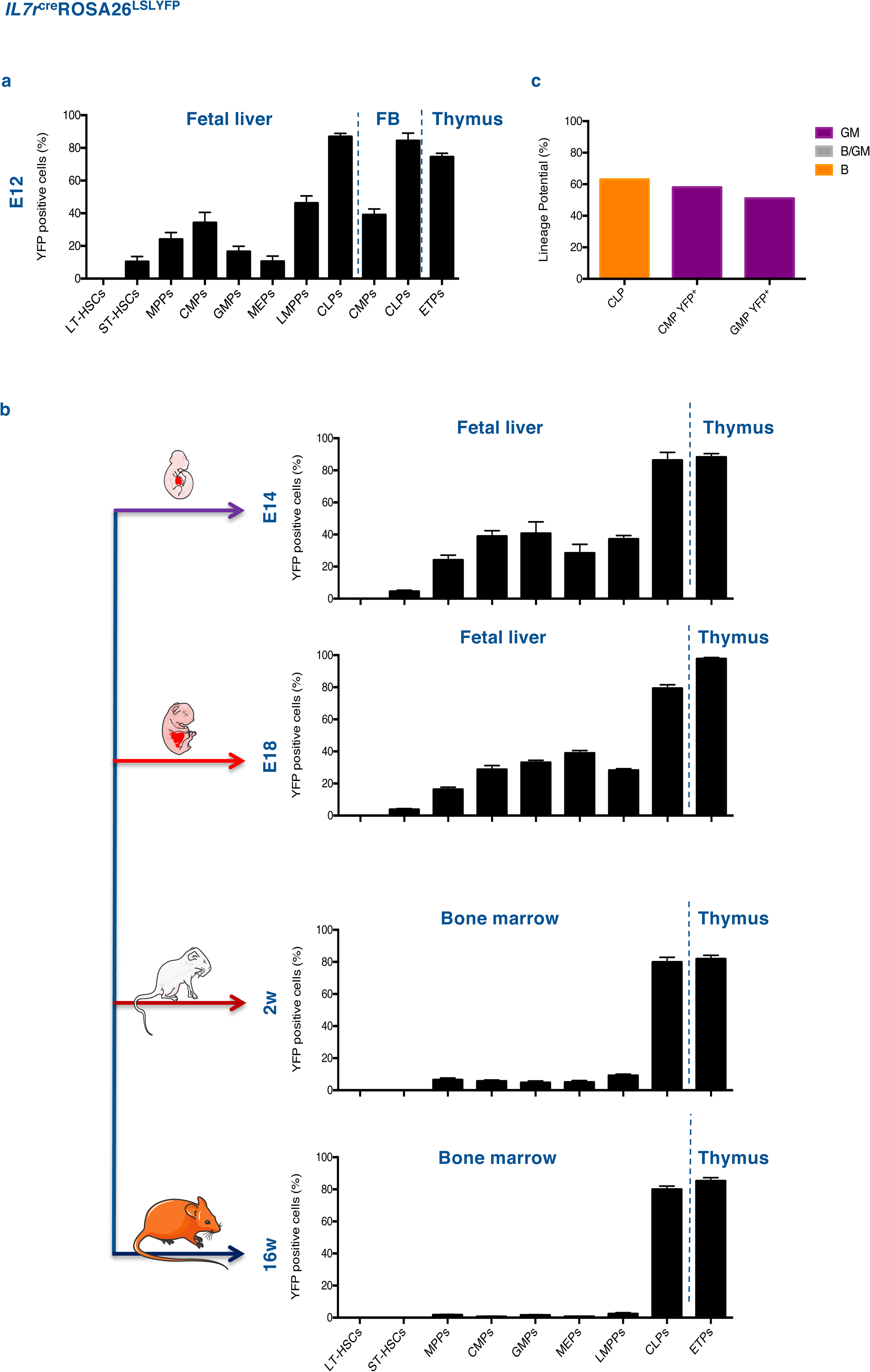
*Il7R⍺^+^* expression in fetal liver multipotent and myeloid progenitors, Related to Figure 4. (a) Frequencies of YFP labelled E12.5 hematopoietic progenitors in the fetal liver, fetal blood (FB) and the thymus of il7r^Cre^-Rosa26^LSLYFP^ embryos. (b) Frequency of wells containing B cells or GM in cultures of single sorted E12.5 fetal liver CLPs, CMP YFP^+^ and GMP YFP^+^ on OP9 cells and analyzed after 12 days of culture (three independent experiments). (c) Frequencies of YFP labelled hematopoietic progenitors analyzed in the fetal liver and fetal thymus of E14.5 and E18.5 embryos or in the bone marrow and thymus of 2 and 16-week old mice. Data are pooled from at least two independent experiments (mean ± s.e.m).

**Figure S5.**
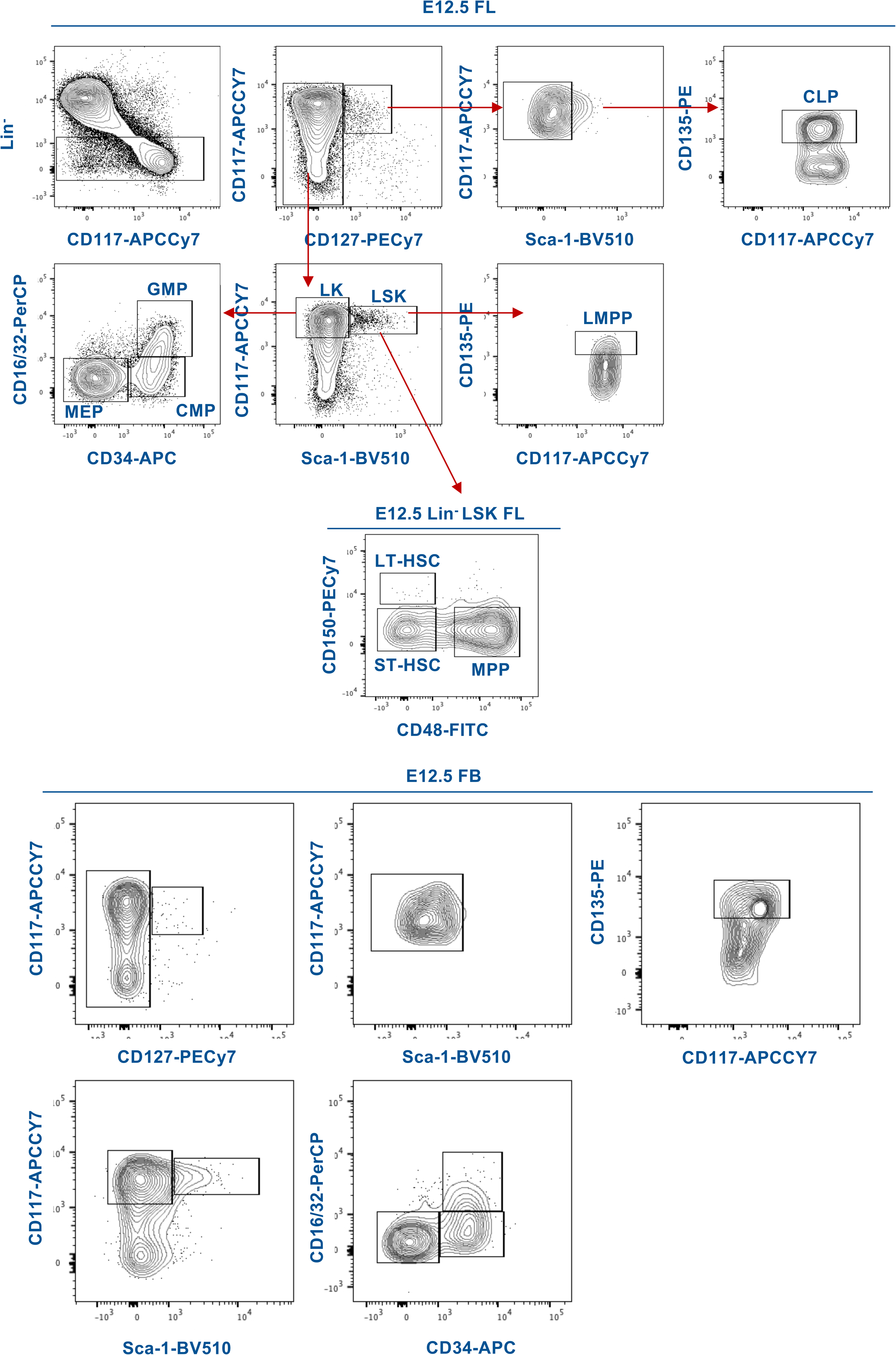
Gating strategy for hematopoietic progenitors, Related to Figure S4.

**Figure S6.**
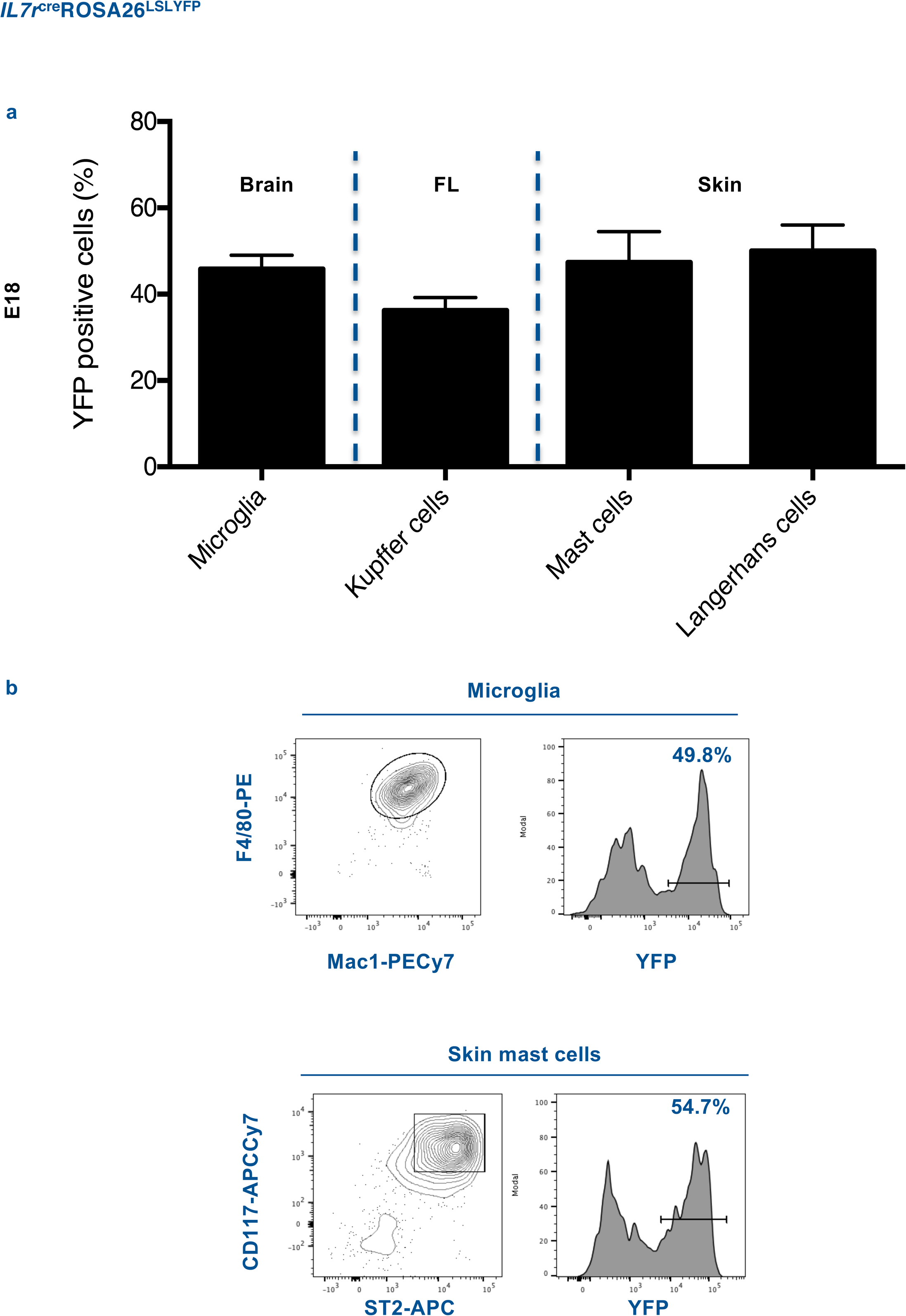
YFP expression in brain, skin and liver resident macrophages and in skin resident mast cells from E18 *Il7r*^cre^ROSA^LSLYFP^ embryos. Related to Figure 4. (a) Frequency of YFP expressing Microglia, liver macrophages, skin macrophages and mast cells of E18 embryos. Data pooled from three independent experiments, data are depicted as mean ± s.e.m. (b) Gating strategy used for the identification of microglia and skin mast cells.

**Figure S7.**
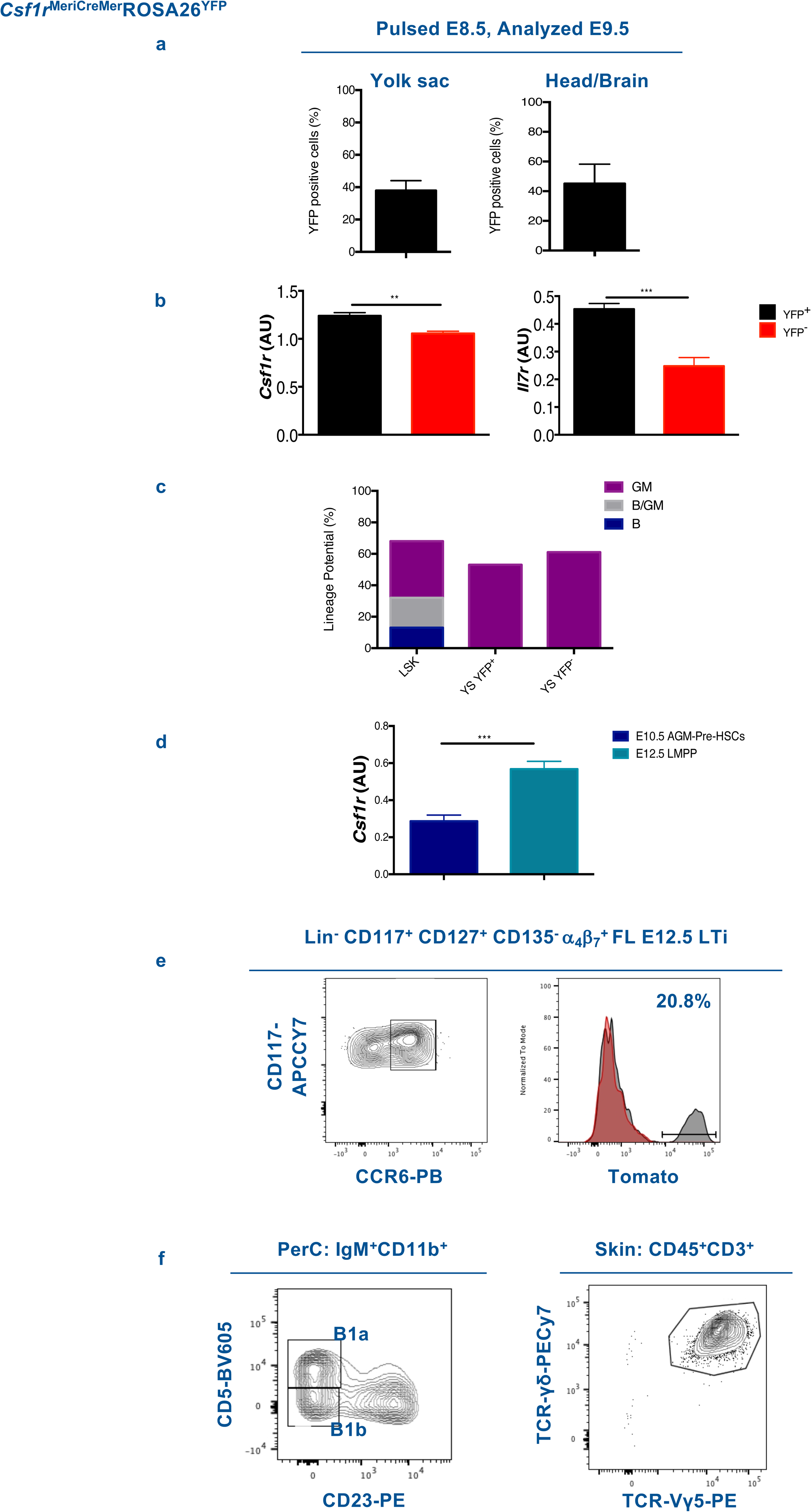
Lineage tracing analysis of *Csf1r*-expressing YS-progenitors. Related to Figure 6. (a) Frequency of YFP+ cells in YS (left histogram) and in the head/Brain (right histogram) of *Csf1r*^MerCreMer^ROSA26^YFP^ E9.5 embryos, pulsed at E8.5. (b) qRT-PCR analysis of *Csf1r* (left graph) and *Il7ra* (right graph) transcripts in E9.5 YS Lin^-^ CD117^+^ CD41^+^ YFP^+^ and YFP^-^ populations. (c) Frequency of B, GM and multiple B/GM cells obtained from single sorted E9.5 YS Lin^-^ CD117^+^ CD41^+^ YFP^+^ and YFP^-^ cells on OP9 stromal cells, cultured with the appropriated cytokines. (d) qRT-PCR analysis of *Csf1r* transcripts in E10.5 AGM-Pre-HSCs (sorted as CD31^+^CD117^+^CD41^+^CD45^low^) and E12.5 FL LMPPs. (e) Flow cytometry gating strategy to identify E12.5 FL lymphoid tissue inducer cells (LTi)(left panel) and frequency of tomato^+^ cells within LTi (right panel). (f) Flow cytometry gating strategy to identify B1 B cells in the peritoneal cavity and Vγ5^+^ T cells in the skin.

**Table 1.**
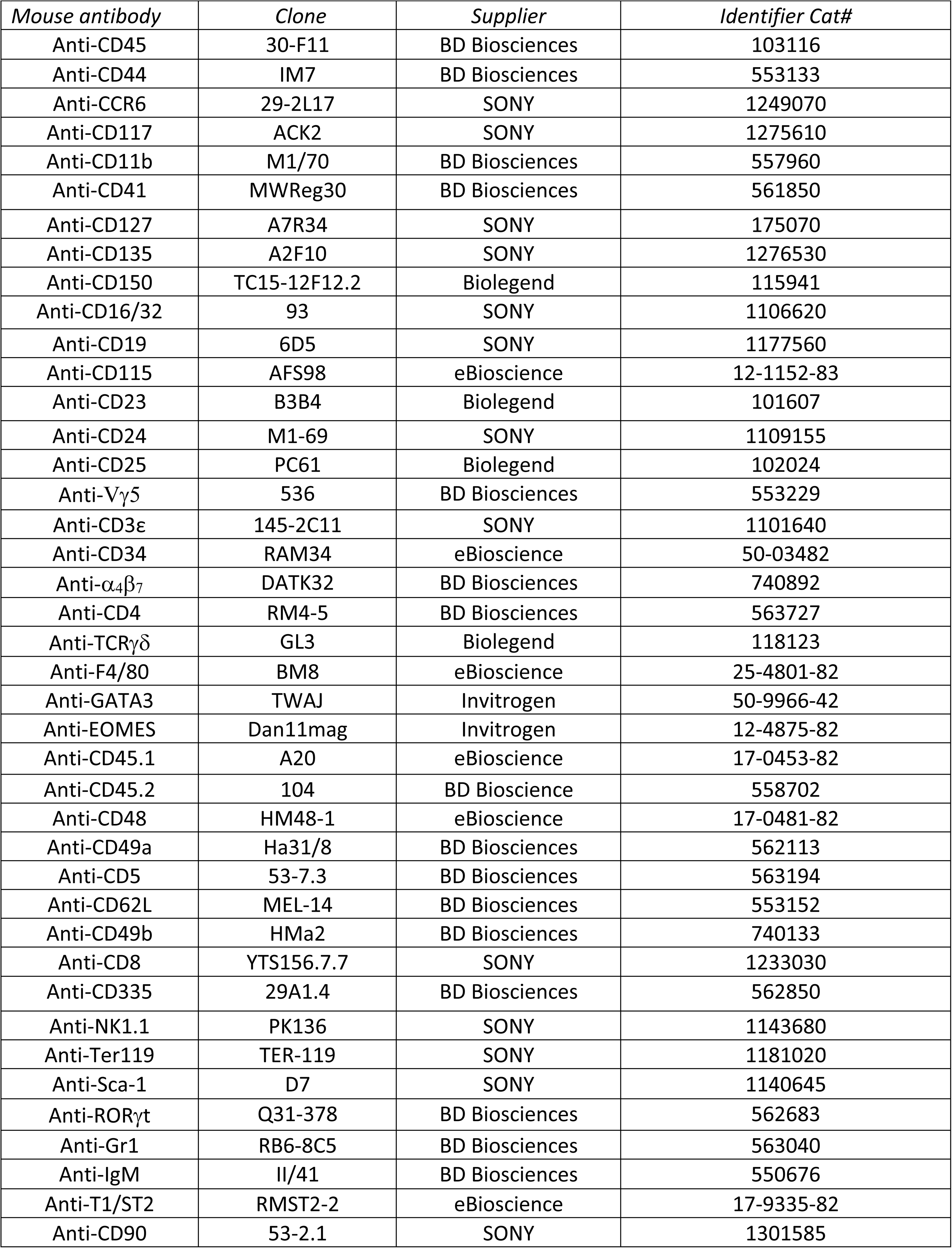
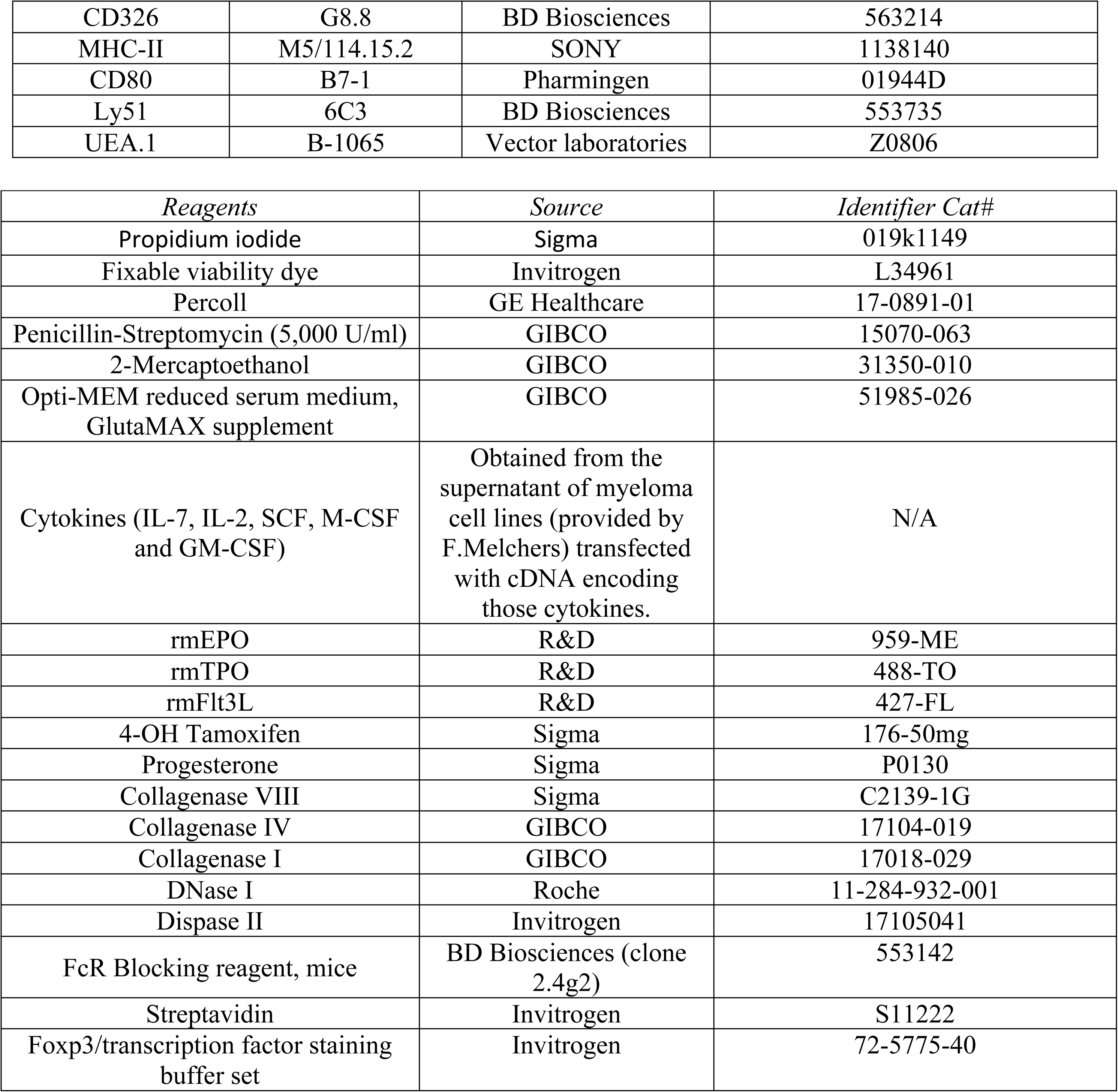
Antibody and reagents list.

**Table 2.**
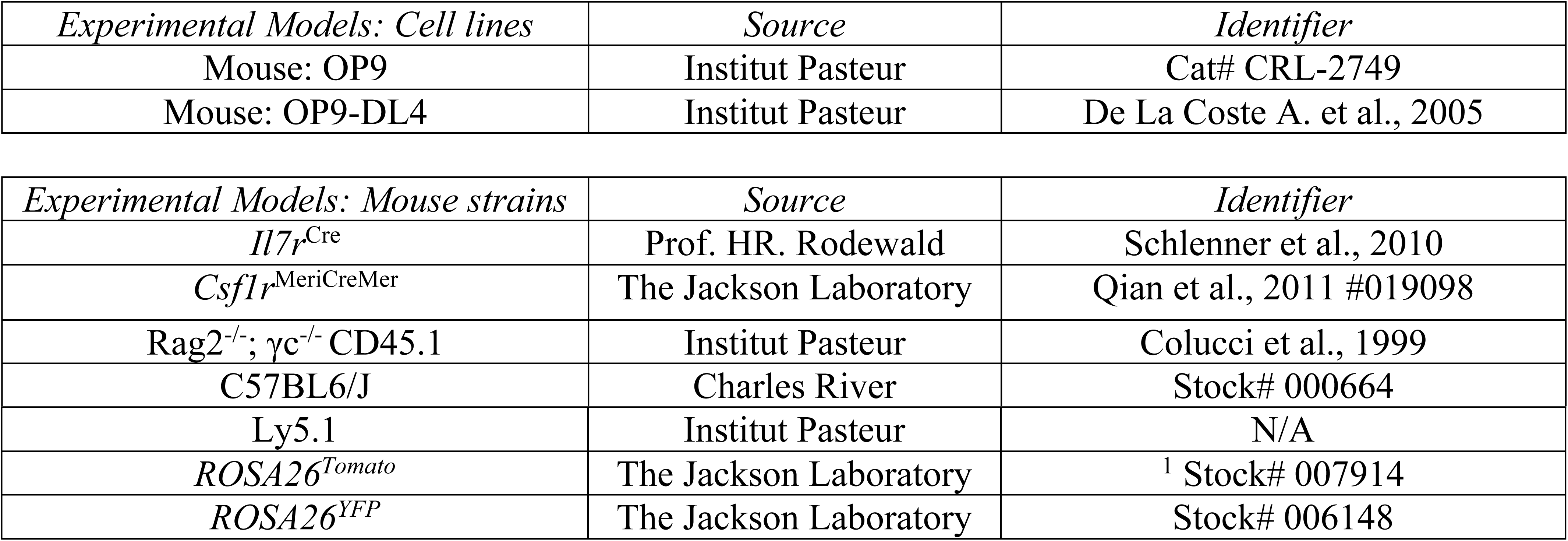
Experimental models.

| *Experimental Models: Cell lines* | *Source* | *Identifier* |
| --- | --- | --- |
| Mouse: OP9 | Institut Pasteur | Cat# CRL-2749 |
| Mouse: OP9-DL4 | Institut Pasteur | De La Coste A. et al., 2005 |

| *Experimental Models: Mouse strains* | *Source* | *Identifier* |
| --- | --- | --- |
| *Il7r<sup>Cre</sup>* | Prof. HR. Rodewald | Schlenner et al., 2010 |
| *Csf1r<sup>MeriCreMer</sup>* | The Jackson Laboratory | Qian et al., 2011 #019098 |
| Rag2<sup>-/-</sup>; γc<sup>-/-</sup> CD45.1 | Institut Pasteur | Colucci et al., 1999 |
| C57BL6/J | Charles River | Stock# 000664 |
| Ly5.1 | Institut Pasteur | N/A |
| *ROSA26<sup>Tomato</sup>* | The Jackson Laboratory | <sup>1</sup> Stock# 007914 |
| *ROSA26<sup>YFP</sup>* | The Jackson Laboratory | Stock# 006148 |

**Table 3.** Biomark genes.

| **Gene** | **Ref** | **Gene** | **Ref** |
| --- | --- | --- | --- |
| *Actb* | *Mm00607939_s1* | *Kit* | *Mm00445212_m1* |
| *Hprt* | *Mm00446968_m1* | *Runx3* | *Mm00450960_m1* |
| *Gapdh* | *Mm03302249_g1* | *Runx1* | *Mm01213405_m1* |
| *Gzma* | *Mm01304452_m1* | *Sox4* | *Mm00486320_s1* |
| *Gzmb* | *Mm00442837_m1* | *Tcf7* | *Mm01173766_m1* |
| *Zbtb16* | *Mm00784709_s1* | *Gata3* | *Mm00484683_m1* |
| *Hes1* | *Mm01342805_m1* | *Tox* | *Mm00455231_m1* |
| *Nfil3* | *Mm00600292_s1* | *Ccr9* | *Mm02620030_m1* |
| *Itga2b* | *Mm00439741_m1* | *Cd32* | *Mm00438875_m1* |
| *Bcl11b* | *Mm00480516_m1* | *Cd34* | *Mm00519283_m1* |
| *Il13* | *Mm00434204_m1* | *Cebpa* | *Mm00514283_s1* |
| *Bcl2* | *Mm_00477631_m1* | *Csf1r* | *Mm00432689_m1* |
| *Igf2* | *Mm00439564_m1* | *Csf2r* | *Mm00438331_g1* |
| *Ccr6* | *Mm99999114_s1* | *Csf3r* | *Mm00432735_m1* |
| *Eomes* | *Mm01351985_m1* | *Ebf1* | *Mm01288946_m1* |
| *Rxrg* | *Mm00436411_m1* | *Epor* | *Mm00438760_m1* |
| *Id2* | *Mm0011781_m1* | *Flk2* | *Mm01210883_m1* |
| *Il1rl1* | *Mm00516117_m1* | *Foxo1* | *Mm00490672_m1* |
| *Tnfrsf11a* | *Mm00437132_m1* | *Gata1* | *Mm00484678_m1* |
| *Tbx21* | *Mm00493445_m1* | *Gata2* | *Mm00492301_m1* |
| *Areg* | *Mm00437583_m1* | *Kdr* | *Mm01222421_m1* |
| *Ncr1* | *Mm01337324_g1* | *Klf1* | *Mm00516096_m1* |
| *Irf8* | *Mm00492567_m1* | *Mpl* | *Mm00440310_m1* |
| *Il7r* | *Mm00434295_m1* | *Myb* | *Mm00501741_m1* |
| *Icos* | *Mm00497600_m1* | *Notch1* | *Mm_00435249_m1* |
| *Rora* | *Mm01173766_m1* | *Pax5* | *Mm00435497_m1* |
| *Rorc* | *Mm01261022_m1* | *Pu.1* | *Mm03048233_m1* |
| *Cd4* | *Mm00442754_m1* | *Rag1* | *Mm01270936_m1* |
| *Cxcr5* | *Mm00432086_m1* | *Rag2* | *Mm00501300_m1* |
| *Il18r1* | *Mm00515178_m1* | *Rbpj* | *Mm01217627_g1* |
| *Il2rb* | *Mm01195267_m1* | *Tal1* | *Mm01187033_m1* |
| *Il1r1* | *Mm00434237_m1* | *Tek* | *Mm00443243_m1* |
| *Ltb* | *Mm00434774_g1* | *Vwf* | *Mm00550376_m1* |
| *Cxcr6* | *Mm02620517_s1* | *Itga4* | *Mm00439770_m1* |
| *Ets1* | *Mm01175819_m1* |  |  |
| *Sell* | *Mm00441291_m1* |  |  |

